# The histone chaperone CAF-1 prevents Rad52-mediated instability of the budding yeast ribosomal DNA during replication-coupled DNA double-strand break repair

**DOI:** 10.1101/2024.03.12.584701

**Authors:** Hajime Futami, Tsugumi Yamaji, Yuko Katayama, Nanase Arata, Takehiko Kobayashi, Mariko Sasaki

## Abstract

DNA replication-coupled chromatin assembly is crucial to maintain genome integrity. Here, we demonstrate that absence of the budding yeast histone chaperone CAF-1 induces chromosomal ribosomal RNA gene (rDNA) copy number changes as well as the production of extrachromosomal rDNA circles in a manner dependent on Fob1 that causes replication fork arrest in the rDNA, the homologous recombination protein Rad52, and its interaction with Proliferating Cell Nuclear Antigen. CAF-1 deficiency enhances transcription from the regulatory promoter E-pro, contributing to rDNA instability, but it also stabilizes the rDNA independently through its regulation of E-pro. Absence of CAF-1 induces end resection of DNA double-strand breaks (DSBs) formed at arrested replication forks, which are repaired in a manner dependent on the Mre11-Rad50-Xrs2 complex. CAF-1 deficiency causes partial defects in lagging strand synthesis coupled to nucleosome spacing in the rDNA. Our findings suggest that CAF-1 suppresses Rad52-mediated rDNA instability during repair of replication-coupled DSBs.

## INTRODUCTION

DNA copy number changes are introduced by deletion and/or amplification of DNA segments on chromosomes or via extrachromosomal circular DNAs derived from chromosomes. These changes can alter gene expression levels, compromise cellular integrity, and drive tumorigenesis and numerous other diseases [reviewed by (1–5)]. Thus, understanding the molecular mechanisms by which chromosomal and extrachromosomal DNA copy number changes occur is important.

The budding yeast genome contains approximately 150 rDNA copies, which are tandemly repeated at a single locus on chr XII (Fig. 1A). Each copy contains 35S and 5S rRNA transcription units, which are separated by intergenic spacers that contain an origin of DNA replication, a cohesin-associated region (CAR), a replication fork barrier (RFB) sequence and a bidirectional RNA polymerase II (RNAP II) promoter, E-pro, which synthesizes noncoding RNA. Once DNA replication is initiated, the DNA replication fork progressing opposite to the 35S rDNA is arrested by the Fob1 protein bound to the RFB site (Fig. 1A) (6–8). The arrested forks are then broken, leading to the formation of a single-ended DSB (9–11). Cells have the mechanisms to maintain their rDNA copy number during DSB repair, and when these mechanisms are deficient, cells undergo changes in chromosomal rDNA copy number and produce extrachromosomal rDNA circles (ERCs) (Fig. 1A) (12).

**Figure 1.**
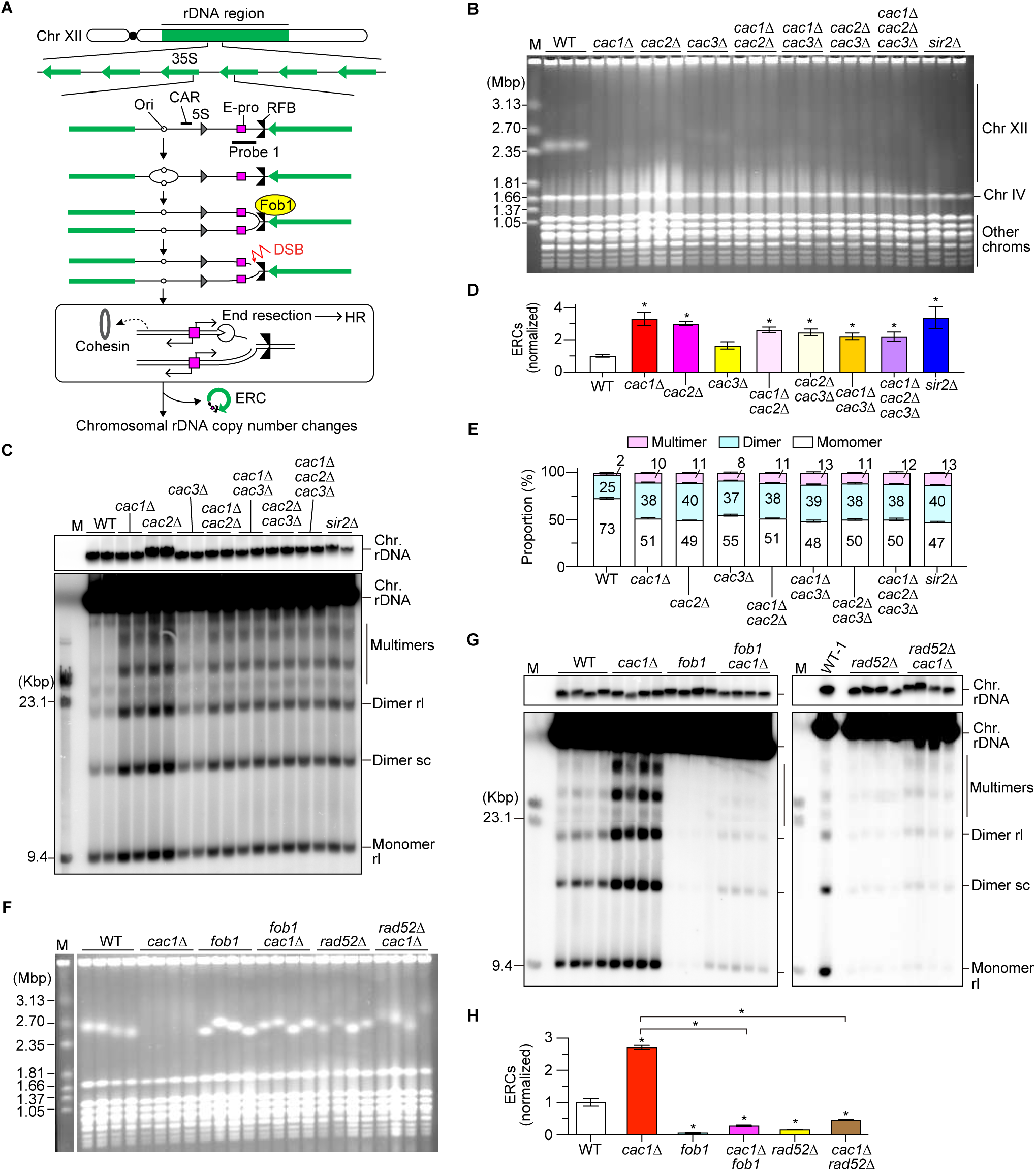
CAF-1 suppresses HR-mediated rDNA instability in response to Fob1-dependent replication fork arrest. **(A)** Replication fork arrest, DSB formation, and rDNA copy number changes in the budding yeast rDNA region. Chr XII carries the rDNA cluster, as indicated by the green bar, where approximately 150 rDNA copies are tandemly arrayed. Each copy contains 35S rRNA (35S), 5S rRNA (5S), an origin of DNA replication (Ori), a cohesin-associated region (CAR), a bidirectional promoter for noncoding RNA, called E-pro, and a replication fork barrier (RFB). The black bar indicates the position of probe 1 used for ERC detection. **(B, F)** PFGE analysis of chr XII size heterogeneity. DNA from three independent clones (B) and four independent clones (F) of the indicated strains was separated by PFGE and stained with ethidium bromide (EtBr). M indicates *Hansenula wingei* chromosomal DNA markers. **(C, G)** ERC detection. DNA was isolated from two independent clones (C) and four independent clones (G) of the indicated strains and separated by agarose gel electrophoresis, followed by Southern blotting with the probe 1, as shown in (A). In (G), DNA from the first WT clone (WT-1) run on the left gel was also loaded on the right gel. Chromosomal rDNA and different ERC forms are indicated. rl and sc indicate relaxed and supercoiled forms of ERCs, respectively. Supercoiled monomers ran off from the gel in the electrophoresis condition used in this experiment. M indicates lambda Hind III DNA markers. The top panel shows short exposure of genomic rDNA signals. **(D, H)** Quantitation of ERCs. ERCs from (C) and Supplementary Figure 1A were quantified in (D). ERCs from (G) were quantified in (H). ERC levels were compared between WT and mutants by one-way ANOVA, followed by Tukey’s multiple comparisons test. Bars show the mean ± s.e.m. Asterisks indicate statistically significant differences between WT and mutant strains (*p* < 0.05). **(E)** Proportion of different ERC forms. In each sample, monomeric, dimeric and multimeric ERCs were quantified and the proportion of each form in total was expressed as proportion of total ERCs.

Homologous recombination (HR) is important for repairing double-ended DSBs formed independently of DNA replication, such as those introduced by endonucleases in the G2/M phase of the mitotic cell cycle and meiotic DSBs [reviewed by (13,14)]. To repair DSBs by HR, 5’-ended strands of the DSB ends are degraded by a process called DSB end resection [reviewed in (15)]. Resected DSBs are subsequently repaired using a homologous sequence on the sister chromatid as a template for Rad52-dependent recombination reactions, which properly restore the original sequence at the DSB site. However, when DSBs are formed in repetitive regions, HR can use a homologous template located at a misaligned position on the sister chromatid or elsewhere in the genome, resulting in chromosome rearrangements [reviewed by (4)].

Cells may use distinct mechanisms to repair DSBs that are formed in the context of DNA replication forks (16). We previously demonstrated that DSB end resection is normally limited for single-ended DSBs formed in the context of arrested replication forks at the rDNA RFB, and these DSBs can be repaired without an essential HR factor, Rad52 (17). We have identified two factors involved in suppression of DSB end resection. First, when cells lack the replisome component Ctf4, DSB end resection is induced, DSB repair requires the MRX complex and Rad52, and subsequent HR-dependent DSB repair results in rDNA hyperamplification (17). The dependency on the MRX complex and HR proteins for DSB repair seen when Ctf4 is absent is similar to the genetic requirements for repair of replication-coupled DSBs introduced by nickases or endonucleases (16,18–22). Secondly, the histone deacetylase Sir2 represses transcription from E-pro (Fig. 1A). When Sir2 is absent, transcription from E-pro is activated, inducing DSB end resection (23). During subsequent DSB repair, HR-mediated reactions are responsible for inducing chromosomal rDNA copy number changes as well as ERC production, but unlike when Ctf4 is absent, cells lacking Sir2 require the MRX complex but not Rad52 to complete DSB repair (23).

Cohesin complexes that keep sister chromatids in close proximity are recruited to the CAR and RFB regions (Fig. 1A) (10,23–25) and are suggested to facilitate the use of the rDNA copy at an aligned position for DSB repair (10,24). The absence of Sir2 and subsequent transcriptional activation lead to the inhibition of cohesin associations with CAR and RFB and rDNA instability (10,23,24). Therefore, the repression of transcription from E-pro is important for maintaining rDNA stability not only by suppressing DSB end resection and subsequent HR-mediated DSB repair but also by ensuring cohesin binding to rDNA (Fig. 1A).

Eukaryotic organisms must faithfully replicate not only their genetic information but also their chromatin states to maintain genome integrity and epigenetic information. During DNA replication, nucleosomes are disassembled to allow the progression of DNA replication forks [reviewed by (26,27)]. Then, the parental histones are recycled and deposited onto the newly replicated DNA. In addition, newly synthesized histones are deposited onto DNA *de novo* by histone chaperones. The histone chaperone Asf1 first binds to the newly synthesized histones H3-H4 and presents them to the Rtt109 acetyltransferase for acetylation at histone H3K56 (28–30). Acetylated H3-H4 is then transferred to other histone chaperones such as CAF-1 and Rtt106. CAF-1, which consists of Cac1, Cac2 and Cac3 in budding yeast, is recruited to the replisome through its interaction with Proliferating Cell Nuclear Antigen (PCNA) and deposits (H3-H4)_2_ tetramers onto the newly synthesized DNA (31–33). These replication-coupled nucleosome assembly factors play important roles in response to DNA damage but the degree to which each factor contributes to different DNA damage repair pathways differs, depending on the source and context of DNA damage (21,34–38). During DNA replication *in vivo*, Okazaki fragment synthesis on the lagging strand is interlinked with the chromatin assembly: Okazaki fragments show a size distribution similar to that of the nucleosome repeat, and Okazaki fragment termini are located near nucleosome midpoints (39). The analysis of bulk DNA synthesized during DNA replication demonstrates that CAF-1 facilitates the generation of properly sized Okazaki fragments by mediating nucleosome deposition behind replication forks (39–41).

Previous studies demonstrate that the CAF-1 complex is required for restricting HR-dependent rDNA copy number loss during DSB repair in *Arabidopsis thaliana* (42–44). Here, we showed that absence of the budding yeast histone chaperone CAF-1 induces not only loss but also gain of chromosomal rDNA copies as well as the production of ERCs in a manner dependent on Fob1 and Rad52. CAF-1 deficiency led to derepression of transcription from E-pro. This transcriptional activation induced rDNA instability in the *caf-1* mutants, but other defects also contributed to this phenotype. Absence of CAF-1 enhanced end resection of replication-coupled DSBs at the RFB, and DSB repair and subsequent ERC production depend on the MRX complex. Mutations that compromise the interaction of CAF-1 with PCNA led to rDNA instability, indicating that CAF-1’s function at the replication forks is important for the maintenance of rDNA stability. Furthermore, CAF-1 deficiency led to a reduction in nucleosome-sized Okazaki fragments in the rDNA. Taken together, these findings suggest that CAF-1 is important for suppressing Rad52-dependent rDNA instability during repair of DSBs formed in the context of arrested replication forks in the rDNA.

## Materials and Methods

### Yeast Strains and Culture Methods

Yeast strains used in this study are derivatives of W303 (*MATa ade2-1 ura3-1 his3-11, 15 trp1-1 leu2-3,112 can1-100*) and are listed in Table S1. Haploid strains lacking genes of interest were constructed by standard one-step gene replacement methods, followed by confirmation of genotypes by PCR. Alternatively, diploids heterozygous for gene deletions of interest were constructed, subjected to sporulation, and haploid clones were isolated by tetrad dissection. Strains carrying plasmids were constructed by standard transformation methods. Yeast cells were grown in YPD media (10 g/L yeast extract, 20 g/L peptone, and 20 g/L glucose). For Fig. 5, cells carrying plasmids were selected in SC-Trp media [6.7 g/L yeast nitrogen base without amino acids (BD, 291940), 1.92 g/L yeast synthetic drop-out medium supplements without tryptophan (Sigma, Y1876), 20 g/L glucose]. Agar plates were prepared similarly, with agar added at 20 g/L.

For PFGE and ERC analyses, five mL of appropriate liquid media was inoculated with cells and grown overnight to saturation at 30°C. Cultures were collected (5 × 10^7^ cells/plug), washed twice with 50 mM EDTA (pH 7.5), and cell pellets were stored at −20°C. For RNA preparation, overnight cultures were diluted in 15 mL of YPAD medium (YPD containing 40 μg/mL adenine sulfate) to OD_600_ = 0.2 and grown until OD_600_ = 0.8. Cultures equivalent to ∼1– 2 × 10^8^ cells were collected, washed twice with ice-cold water, snap-frozen in liquid nitrogen, and stored at −80°C. For protein extracts, overnight cultures were diluted in 15 mL to OD_600_ =0.2 and grown until OD_600_ = 0.8. Cultures equivalent to ∼0.5–1 × 10^8^ cells were collected, washed twice with ice-cold water, and stored at −80°C. For chromatin immunoprecipitation, overnight cultures of MSY426, HFY106, and HFY122 were diluted in 100 mL of YPD medium to an OD_600_ = 0.2 and grown until OD_600_ = 0.8. Cells were immediately subjected to crosslinking as described below. Gal-pro WT/*cac1*Δ/*sir2*Δ strains (HF70, 73, 76) were first grown in 5 mL YPD media overnight at 30°C. Cells were inoculated at ∼4–5 × 10^4^ cells/mL into 15 mL of YP media containing 20% raffinose and grown for >24 hrs at 30°C. Cells were then inoculated at 2 × 10^6^ cells/mL into two sets each of 7 mL of YP media containing either 20% raffinose or 20% galactose (one culture for RNA preparation and the other for agarose plug preparation). To prepare RNA preparation, cultures were grown until OD_600_ = 0.8, collected, washed twice with ice-cold water, snap-frozen in liquid nitrogen, and stored at −80°C.

For DSB analyses, time course experiments were performed with MSY360, 937, 1231 1261, and 1262, as described previously (17,45). *GAL-FOB1* strains were grown overnight in YPD media at 30°C. Cells were inoculated into ∼200 mL of YP media supplemented with 40 μg/mL adenine sulfate, 2% raffinose, and one drop of antifoam 204 (Sigma) at a density to reach ∼1 × 10^7^ cells/mL the following morning and grown overnight at 23°C. To arrest cells in G1 phase, α-factor (synthesized by BEX Co. Ltd. [Tokyo, Japan]) was added to cultures at a final concentration of 20 nM. Cells were incubated at 23°C for 3–4 hrs until >80% of the cells were arrested in G1. Then, 20% galactose was added at a final concentration of 2% to induce *FOB1* expression and the cultures were further incubated for 30 min at 23°C. Cells were collected by centrifugation for 5 min at 4°C at 4,500 × *g* and washed twice with YP medium containing 40 μg/mL adenine sulfate and 2% galactose. Cell pellets were resuspended at ∼1 × 10^7^ cells/mL in fresh YP medium containing 40 μg/mL adenine sulfate, 2% galactose, 0.075 mg/mL pronase E (Sigma), and one drop of antifoam 204 (Sigma), then incubated at 23°C. At various time points, 10 mL culture aliquots equivalent to 1 × 10^8^ cells were collected, immediately treated with 1/100 volume of 10% sodium azide, collected, washed twice with 50 mM EDTA pH 7.5, and stored at −20°C. Additional 1 mL culture aliquots were collected for flow cytometry analysis.

For analysis of Okazaki fragments, MSY613, which was derived from yIW310 that is a kind gift of Iestyn Whitehouse, MSY1599, MSY1604, MSY1605 were grown overnight in YPD media at 30°C. Overnight cultures were diluted in 200 mL of YPD medium to OD_600_ =0.1 and grown to OD_600_ = 0.4. Each culture was split into two flasks. Doxycycline (100 mg/mL) dissolved in DMSO was added to one culture at a final concentration of 40 μg/mL, while an equal volume of DMSO was added to the other culture. After 2.5 hr incubation at 30°C, cells were collected and washed twice with 50 mM EDTA (pH 8.0), and stored at −20°C.

### Plasmid construction

Plasmids carrying *POL30* (pBL230-0) and mutant alleles *pol30-6* (pBL230-6), *pol30-8* (pBL230-8), and *pol30-79* (pBL230-79) were kind gifts from Peter Burgers (46,47). As these plasmids carry *TRP1* as a selection marker, cells transformed by these plasmids were selected in SC-trp media. YCp22-CAC1 plasmid was constructed by Gibson assembly as follows. The genomic region containing the *CAC1* open reading frame with a few hundred bp upstream and downstream regions were amplified by PCR from the *S. cerevisiae* WT strain of the W303 background, using primers 5’-TGCATGCCTGCAGAGATCTACATCGTTAAACCAAAAG and 5’-GAGCTCGGTACCTTGCATGCGAGAAGTGCTG. Linearized YCplac22 was generated by PCR from YCplac22 with primers 5’- GATGTAGATCTCTGCAGGCATGCAAGCTTG and 5’-TCTCGCATGCAAGGTACCGAGCTCGAATTC. These PCR fragments had short homology at each end and were fused using Gibson Assembly (New England Biolabs) according to the manufacturer’s instructions. YCp22-cac1-20 plasmid carrying the *cac1-20* mutation was constructed using the PrimeSTAR Mutagenesis Basal Kit (TaKaRa) with YCp22-CAC1 as a template and primers 5’-GTAACGCCGGTAAAAAACTAAGCGATTCTAATA and 5’- TTTTACCGGCGTTACCAATACGGGATTGTGCTC, according to the manufacturer’s instructions.

YCp22-CAC1-9myc and YCp22-cac1-20-9myc plasmids were constructed by inserting the sequence that encodes nine consecutive myc epitope tag prior to the stop codon of *CAC1* and *cac1-20* as follows. The 9myc sequence was amplified by PCR using pYM19 (48) and primers 5’-CCAACCCCGTCTTTGCGTACGCTGCAGGTCGAC and 5’- GATCCGTTCAAGTTAGCTAGTGGATCCGTTC. YCp22-CAC1 and YCp22-cac1-20 were linearized with the primers 5’-CAAAGACGGGGTTGGC and 5’- TAACTTGAACGGATCTTTAGTATATAGAA. These PCR fragments had short homology at each end and were fused using In-Fusion Snap Assembly Master Mix (TaKaRa Bio Inc., Japan), according to the manufacturer’s instructions. Plasmid DNA was sequenced to confirm that it had the proper mutation or insertion.

### Genomic DNA preparation

For PFGE, ERC, and DSB gel analyses, genomic DNA was prepared in low melting temperature agarose plugs as described previously (17,45). Briefly, collected cells were resuspended in 50 mM EDTA pH 7.5 (33 μL per 5 × 10^7^ cells) and incubated at 42°C. For each plug, 33 μL cell suspension was mixed with 66 μL of solution 1 (0.83% low-melting-point agarose SeaPlaque GTG (Lonza), 170 mM sorbitol, 17 mM sodium citrate, 10 mM EDTA (pH 7.5), 0.85% β-mercaptoethanol, and 0.17 mg/mL Zymolyase 100 T (Nacalai)), poured into a plug mold (Bio-Rad) and placed at 4°C until the agarose was solidified. Plugs were transferred to a 2 mL tube containing solution 2 (450 mM EDTA pH 7.5, 10 mM Tris-HCl pH 7.5, 7.5% β-mercaptoethanol and 10 μg/mL RNaseA (Macherey-Nagel)) and incubated for 1–1.25 hrs at 37°C. Plugs were then incubated overnight at 50°C in solution 3 (250 mM EDTA pH 7.5, 10 mM Tris-HCl pH 7.5, 1% sodium dodecyl sulfate (SDS) and 1 mg/mL proteinase K (Nacalai)). Plugs were washed four times with 50 mM EDTA (pH 7.5) and stored at 4°C in 50 mM EDTA (pH 7.5).

For analysis of Okazaki fragments, genomic DNA was isolated as described previously (39), with slight modifications. Briefly, cells were washed twice with lysis buffer (50 mM Tris-HCl [pH 8.0], 50 mM EDTA [pH 8.0], 100 mM NaCl, and 1.5% SDS). Cells were then resuspended in 900 μL of SCE solution (1 M sorbitol, 0.1 M sodium citrate, 0.06 M EDTA [pH 7.5]) supplemented with 35 μL of β-mercaptoethanol and 100 μL of 50 mg/mL Zymolyase and mixed by vortexing. After incubating the cell suspension for 3 min at room temperature, spheroplasts were collected by centrifugation for 5 min at 4,500 × *g* and rinsed with 1 mL of SCE and collected by centrifugation for 5 min at 4,500 × *g*. Spheroplasts were resuspended in 500 μL of lysis buffer (50 mM EDTA (pH 8.0), 50 mM Tris-HCl (pH 8.0), 0.5% sodium dodecyl sulfate, and 200 μg/mL proteinase K) and incubated for 2 hrs at 42°C. Proteins were precipitated by adding 200 μL of 5 M potassium acetate, incubating for 5 min on ice, and centrifuging at 15,000 rpm for 15 min at 4°C. DNA in the supernatant was precipitated with isopropanol and resuspended in 250 μL of STE (10 mM Tris-HCl [pH 8.0], 1 mM EDTA [pH 8.0], 100 mM NaCl containing 80 μg/mL RNaseA [Macherey-Nagel]). RNA was digested by incubation for 1 hr at 37°C. DNA was ethanol precipitated, suspended in 50 μL of TE, and quantified using a Qubit Fluorometer.

## PFGE

PFGE was performed as described previously (17,45). Briefly, one-third of a plug was placed on a tooth of the comb, alongside a piece of *Hansenula wingei* chromosomal DNA markers (Bio-Rad). The comb was placed in a gel tray, and 1.0% agarose solution (Pulsed Field Certified Agarose, Bio-Rad) in 0.5× TBE (44.5 mM Tris base, 44.5 mM boric acid and 1 mM EDTA pH 8.0) was poured. PFGE was performed on a Bio-Rad CHEF Mapper XA Chiller System in 2.2 L of 0.5× TBE under the following conditions: 3.0 V/cm for 68 hrs at 14°C, 120° included angle, with initial and final switch times of 300 s and 900 s, respectively. After electrophoresis, DNA was stained with 0.5 μg/mL ethidium bromide (EtBr) for 30 min, washed with dH_2_O for 30 min, and then photographed.

### Southern blotting

#### Agarose gel electrophoresis

##### ERC assay

The ERC assay was performed as described previously (23,45). Half of the agarose plug was placed on a tooth of the comb. After setting the comb in the gel tray (15 × 25 cm), 300 mL of 0.4% agarose (SeaKem LE Agarose, Lonza) in 1× TAE (40 mM Tris base, 20 mM acetic acid, and 1 mM EDTA pH 8.0) was poured into the tray and allowed to solidify. Lambda HindIII DNA marker (600 ng) was loaded in an empty lane. Electrophoresis was performed on a Sub-cell GT electrophoresis system (Bio-Rad) in 1.45 L of 1× TAE at 1.0 V/cm for ∼20–48 hrs at 4°C with buffer circulation. The buffer was changed every ∼24 hrs. DNA was stained with 0.5 μg/mL ethidium bromide (EtBr) for 30 min and then photographed.

##### DSB assay

The DSB assay was performed as described previously (17,45). One-third of an agarose plug was cut and placed in a 2 mL tube. The plug was equilibrated four times in 1 mL of 1× TE (10 mM Tris base, pH 7.5 and 1 mM EDTA, pH 8.0) by rotating the tube for 15 min at room temperature. The plug was then equilibrated twice in 1 mL of 1× NEBuffer 3.1 (New England Biolabs) by rotating the tube for 30 min at room temperature. After discarding the buffer, the plug was incubated in 160 μL of 1× NEBuffer 3.1 buffer containing 160 units of Bgl II (New England Biolabs) overnight at 37°C. The plug was placed on a tooth of the comb which was set into the gel tray (15 × 25 cm), into which 0.7% agarose solution in 1× TBE was poured; after the gel was solidify, 600 ng of lambda BstEⅡ DNA markers were loaded in an empty lane. Electrophoresis was performed on a Sub-cell GT electrophoresis system (Bio-Rad) in 1.45 L of 1× TBE at 2.0 V/cm for 22 hrs at room temperature with buffer circulation. After electrophoresis, DNA was stained with 0.5 μg/mL EtBr for 30 min and then photographed.

##### Okazaki fragment assay

DNA samples were prepared in 20 μL containing 500 ng of genomic DNA, 2 μL of 10× alkaline buffer (0.5 M NaOH, 10 mM EDTA [pH 8.0]), 2 μL of 0.1% bromophenol blue, and 1 μL of 50% glycerol, which were heat-denatured for 5 min at 70°C, followed by rapid chilling on ice. A total of 500 ng of 100 bp DNA ladder was similarly denatured. The agarose solution was prepared by dissolving 2.5 g of Seakem LE agarose (Lonza) in 225 mL of Milli-Q water by microwave, followed by cooling to 50°C. Then, 25 mL of 10× alkaline buffer was added to the agarose solution, mixed, poured into a gel tray (15 × 20 cm) in a cold room, and left until the gel solidified. The comb was removed, and 1.45 L of 1× alkaline buffer was added. DNA samples were loaded. The gel was run overnight for 4 hrs at 4°C at 80 V.

#### DNA transfer

After agarose gel electrophoresis, DNA was transferred to Hybond-XL (GE Healthcare) by the standard capillary transfer method. Alternatively, DNA was vacuum-transferred to Hybond-XL (GE Healthcare) or Nytran SPC (Cytiva) using VacuGene XL (GE Healthrecare). Vacuum was applied at 55 cm Hg for 30–45 min with freshly prepared 0.25 N HCl, for 20–30 min with denaturation solution (0.5 N NaOH, 1.5 M NaCl), and for 1 hr 40 min–2 hrs with transfer buffer (0.25 N NaOH, 1.5 M NaCl). After transfer, DNA was fixed to the membrane by soaking the membrane in 300 mL of freshly prepared 0.4 N NaOH for 10 min with gentle shaking, followed by rinsing the membrane with 2× SSC for 10 min. The membrane was subsequently dried and stored at 4°C.

#### Probe preparation

Probes were prepared as described previously (17,45). Double-stranded DNA fragments were amplified by PCR. Probe 1 used for ERC and Okazaki fragment analyses was amplified with the primers 5’-CATTTCCTATAGTTAACAGGACATGCC and 5’- AATTCGCACTATCCAGCTGCACTC, and probe 4 used for DSB analysis was amplified with the primers 5’-ACGAACGACAAGCCTACTCG and 5’-AAAAGGTGCGGAAATGGCTG. A portion of the PCR products was gel-purified and used as templates for a second round of PCR with the same primers. The PCR products were then gel-purified. Fifty ng of purified PCR products was used for random priming reactions in the presence of the radiolabeled nucleotide, [α-^32^P]-dCTP (3,000 Ci/mmol, 10 mCi/mL, Perkin Elmer), using Random Primer DNA Labeling Kit (TaKaRa), according to the manufacturer’s instructions. For Fig. 4G and 4I, 50 ng of gel-purified PCR products were labeled with digoxigenin (DIG)-11-dUTP using DIG-High Prime (Roche), according to the manufacturer’s instructions. Unincorporated nucleotides were removed using ProbeQuant G-50 Micro Columns (GE Healthcare). Radiolabeled probes were heat-denatured for 5 min at 100°C, immediately prior to hybridization to the membrane.

#### Hybridization

Southern hybridization was performed as described previously (17,23,45). The membrane was prewetted with 0.5 M phosphate buffer pH 7.2 and prehybridized for 1 hr at 65°C with 25 mL of hybridization buffer (1% bovine serum albumin, 0.5 M phosphate buffer, pH 7.2, 7% SDS, 1 mM EDTA pH 8.0). After discarding the buffer, the membrane was hybridized overnight at 65°C with 25 mL of hybridization buffer containing heat-denatured probe. The membrane was washed four times for 15 min at 65°C with wash buffer (40 mM phosphate buffer pH 7.2, 1% SDS, 1 mM EDTA pH 8.0) and exposed to a phosphor screen.

Non-radioactive hybridization in Fig. 4G and 4I was performed using DIG-High Prime DNA Labeling and Detection Starter Kit II, CPD-*Star*, and DIG Wash and Block Buffer Set (Roche) according to the manufacturer’s instructions. Briefly, membranes were prewetted with 2× SSC and prehybridized for 1 hr at 42°C in DIG Easy Hyb buffer. After the buffer was discarded, membranes were hybridized overnight at 42°C with DIG Easy Hyb buffer containing heat-denatured probe. Membranes were washed twice for 15 min at 65°C with 2× SSC, 0.1% SDS and then twice for 15 min at 65°C with 0.5× SSC, 0.1% SDS. After a 5 min wash at room temperature with Washing buffer, membranes were blocked for 30 min at room temperature with Blocking solution, and Blocking solution was discarded. Membranes were incubated for 30 min at room temperature with Anti-Digoxigenin AP diluted by 20,000 fold in Blocking solution. Membranes were washed twice for 15 min with Washing solution and treated for 5 min with Detection buffer. CDP *Star* was added onto the membrane drop-wise, followed by incubation for 5 min. Then, chemiluminescent signals were detected on Bio-Rad ChemiDoc MP.

#### Image analysis

##### ERC assay

Radioactive signals were detected using a Typhoon FLA 7000 or 9000 (GE Healthcare). The chromosomal rDNA signal was used for normalization of the ERC signals. Thus, the membranes hybridized with radioactive probes were exposed to the phosphor screen for a short time and scanned before signal saturation. Then, membranes were re-exposed to a phosphor screen for several days to achieve optimal signal-to-noise ratios for ERCs. For detection of DIG-labeled probes, chemiluminescent signals were detected after short exposure before signal saturation. Subsequently, membranes were exposed for longer time to achieve optimal signal-to-noise ratio for ERCs. ERC bands and chromosomal rDNA band were quantified from images after long and short exposures, respectively, using FUJIFILM Multi Gauge version 2.0 software (Fujifilm) or Amersham ImageQuant TL ver. 10.2 (Cytiva). ERC levels were calculated as the ratio of ERC signal intensities to chromosomal rDNA signals. When the levels of ERCs were compared between samples loaded on different gels, control DNA plugs were included on each gel for normalization.

##### DSB assay

Membranes were exposed to a phosphor screen for 1–4 days until signals of linear molecules, but not arrested forks, DSBs, or resected DSB signals, reached saturation. The frequency of DSBs and resected DSBs was determined as described previously (17,45). Briefly, in each lane, signal intensities of arrested forks, DSBs, resected DSBs, and the terminal fragment containing the telomere-proximal rDNA repeat and its adjacent non-rDNA fragment were quantified using FUJIFILM Multi Gauge version 2.0 software (Fujifilm) or Amersham ImageQuant TL ver. 10.2 (Cytiva). All signals were normalized to the terminal fragment, and the value at t = 0 min was then subtracted from those at other time points. The DSB frequency was calculated as the ratio of maximum DSB level to maximum arrested fork level in the same time course. The frequency of DSB end resection was determined by dividing the maximum level of resected DSBs by the maximum level of DSBs in the same time course.

##### Okazaki fragment assay

Membranes were exposed to a phosphor screen for several days. Lane profiles of signal intensity of each lane were analyzed using ImageJ (NIH).

### Yeast RNA preparation

RNA was prepared as described previously (49), with slight modifications. Collected cells were resuspended in 400 μL of TES (10 mM Tris-HCl pH 7.5, 10 mM EDTA pH 7.5, 0.5% SDS) and 400 μL of acidic phenol by vortexing for approximately 10 s. Cells were incubated at 65°C for 1 hr with occasional vortexing every 15 min. Cell suspensions were then incubated for 5 min on ice and centrifuged at 20,000 × *g* for 10 min at 4°C. The aqueous phase was transferred to a new tube and mixed with an equal volume of acidic phenol by vortexing for 10 s. After 5 min incubation on ice and centrifugation at 20,000 × *g* for 10 min at 4°C, the aqueous phase was transferred to a new tube. RNA was precipitated overnight at −20°C by adding 1/10 volume of 3 M sodium acetate (pH 5.3) and 2.5 volumes of 100% ethanol. The precipitate was collected by centrifugation at 20,000 × *g* for 10 min at 4°C and washed with 70% ethanol. RNA pellets were resuspended in 30 μL of dH_2_O treated with 0.1% diethylpyrocarbonate (DEPC). RNA concentration was determined using a NanoDrop ND-1000 spectrophotometer (Thermo Fisher Scientific). Samples were stored at −80°C.

### Northern blotting

Total RNA (30 μg) was brought up to 7 μL with DEPC-treated dH_2_O and mixed with 17 μL of RNA sample buffer (396 μL of deionized formamide, 120 μL of 10× MOPS buffer [0.2 M MOPS, 50 mM sodium acetate pH 5.2, 10 mM EDTA pH 7.5 in DEPC-treated dH_2_O], and 162 μL of 37% formaldehyde). A total of 1.8 μg of DynaMarker RNA High Markers (BioDynamics Laboratory) were brought up to 4.8 μL with DEPC-treated dH_2_O and mixed with 11.2 μL of RNA sample buffer. Samples were heated at 65°C for 20 min and rapidly chilled on ice. To the samples, 6 μL of 6× Gel Loading Dye (B7025S, New England Biolabs) and 1.5 μL of 2.5 mg/mL EtBr were added. To the size markers, 4 μL of 6× Gel Loading Dye (B7025S, New England Biolabs) and 1 μL of 2.5 mg/mL EtBr were added.

Two 1% agarose gels were prepared by dissolving 1 g of agarose in 73 mL of DEPC-treated dH_2_O. After cooling to 60°C, 17 mL of 37% formaldehyde and 10 mL of 10× MOPS buffer were added. The solution was poured into a gel tray (13 × 12 cm) and allowed to set. Agarose gel electrophoresis was performed on a Mupid EX system (Takara) in 400 mL of 1× MOPS buffer at 20 V for 20 min and then at 100 V until the bromophenol blue dye migrated approximately 2/3 of the gel. Gels were photographed and RNA was transferred to Hybond-N+ (GE Healthcare) by standard capillary transfer.

Strand-specific probes were prepared from double-stranded DNA fragments, amplified by PCR and gel-purified. The PCR primers used for probe 2 (against IGS1-R, IGS2-R), 5’- TCGCCAACCATTCCATATCT and 5’-CGATGAGGATGATAGTGTGTAAGA; for probe 3 (against IGS1-F) were 5’-AGGGAAATGGAGGGAAGAGA and 5’-TCTTGGCTTCCTATGCTAAATCC, and for detecting *TSA1*, 5′- CAAGTTCAAAAGCAAGCTCC and 5′-ACCAATCTCAAGGCTTCGTC. Strand-specific probes were then prepared by linear PCR in a final volume of 20 μL containing 0.2 mM dATP, 0.2 mM dTTP, 0.2 mM dGTP, 50 μL [α-^32^P]-dCTP (3,000 Ci/mmol, 10 mCi/mL, Perkin Elmer), 1.25 units ExTaq (TaKaRa), 1× ExTaq buffer, 50 ng of PCR product as a template, and 10 μM primer (5’-AGTTCCAGAGAGGCAGCGTA for probe 2, 5’-CATTATGCTCATTGGGTTGC for probe 3). PCR was initiated by a denaturation step at 94°C for 3 min, followed by 35 cycles of amplification (96°C for 20 s, 51°C for 20 s, and 72°C for 30 s), and a final step at 72°C for 3 min. Unincorporated nucleotides were removed using ProbeQuant G-50 Micro Columns (GE Healthcare). Radiolabeled probes were heat denatured by incubating for 5 min at 100°C, immediately prior to hybridization to the membrane.

Membrane was incubated with 10 mL of ULTRAhyb Ultrasensitive Hybridization Buffer (Thermo Fisher) at 42°C for 1 h. The heat-denatured probe was incubated with the membrane overnight at 42°C. The membrane was rinsed twice with 2× SSC, washed for 15 min at 42°C twice with wash buffer 1 (2× SSC, 0.1% SDS), and washed for 15 min at 42°C twice with wash buffer 2 (0.1× SSC, 0.1% SDS). The membrane was exposed to phosphor screens for several days, and radioactive signals were detected using a Typhoon FLA 7000 (GE Healthcare). Probes were stripped by incubating the membrane with boiled 0.1% SDS by shaking for ∼30 min, rinsed with 2× SSC, and rehybridized with the *TSA1* probe that was prepared as described above. Signals of IGS-F, IGS1-R, IGS2-R, and *TSA1* were quantified using FUJIFILM Multi Gauge version 2.0 software (Fujifilm). The levels of IGS transcripts were normalized to the *TSA1* signal.

### Preparation of protein extracts

Protein extracts were prepared from yeast cells, as described previously with slight modifications (49). Cells were resuspended in 500 μL of ice-cold dH_2_O and 75 μL of Alkali-2ME solution (1.295 N NaOH and 7.5% v/v 2-mercaptoethanol) by vortexing and incubated on ice for 10 min. Cell suspensions were mixed with 75 μL of 50% w/v trichloroacetic Acid by vortexing and incubated on ice for 10 min. After centrifugation at 10,000 × *g* for 5 min at 4°C, the supernatant was completely removed and pellets were resuspended in 75 μL of buffer (800 μL of 200 mM Tris-HCl pH 8.8 and 200 μL of 5x SDS-loading buffer [250 mM Tris-HCl, pH 6.8, 10% SDS, 0.1% bromophenol blue, 25% glycerol, 25% β-mercaptoethanol]). Protein concentration was quantified using a Bradford assay (Bio-Rad). Protein samples were stored at −20°C.

### Western blotting

Protein samples (30 μg) were mixed with 1× SDS-loading buffer to a final volume of 10 μL. Protein samples were heated for 5 min at 65°C, and centrifuged at 20,000 × *g* for 5 min at 4°C. Supernatants were loaded and separated on 5–20% pre-made Bullet PAGE Plus Precast Gel (Nacalai Tesque) in 400 mL of 1x SDS-PAGE running buffer (25 mM Tris, 192 mM Glycine, 0.1% SDS). Proteins were transferred to Immobilon-P PVDF membrane (Merck-Millipore) in 1× transfer buffer (25 mM Tris, 192 mM glycine, 10% methanol) for 60 min at 100 V at 4°C using a Mini Trans-Blot cell (Bio-Rad). After transfer, the membrane was blocked in 5% Skim Milk (Wako) in TBS-T (1x TBS, 0.1% Tween20) for 1 hr at room temperature. The membrane was incubated overnight at 4°C with anti-myc antibody (9E10) (Santa Cruz, sc-40) diluted by 1,000-fold in 1% skim milk in TBST. The membrane was washed three times for 5 min with TBS-T. The membrane was further incubated with anti-mouse IgG-HRP (GE Healthcare, NA931VS) diluted by 10,000-fold in 1% milk in TBST for 1 hr at room temperature. The membrane was washed three times for 5 min with TBS-T. The membrane was incubated with ECL Prime (GE Healthcare) and chemiluminescent signal was detected on Bio-Rad ChemiDoc MP. Antibodies were stripped by incubating for 30 min at room temperature in Western BLoT Stripping Buffer (TaKaRa Bio). From the same membrane, tubulin was detected as a loading control. The membrane was blocked in 5% Skim Milk in TBS-T for 1 hr at room temperature. The membrane was incubated overnight at 4°C with anti-tubulin-HRP (YL1/2) (BIO-RAD, BCA77P) diluted by 1,000-fold in 1% milk in TBS-T. After the membrane was washed three times for 5 min with TBS-T, the membrane was incubated with ECL Prime (GE Healthcare) and chemiluminescent signal was detected on BIO-RAD ChemiDoc MP. Protein signals were quantified using Image Lab software (BIO-RAD).

### Chromatin immunoprecipitation-quantitative PCR analysis

Thirty-five milliliters of cultures were transferred to 50 mL conical tubes and crosslinked with 972 μL of 37% formaldehyde (1% final concentration) by shaking for 30 min at room temperature on a reciprocal shaker at 60 r/min. The reaction was quenched with glycine (a final concentration of 125 mM) by shaking for 5 min at room temperature on a reciprocal shaker at 60 r/min. Cells were collected by centrifugation at 2,380 × *g* for 3 min at 4°C, washed twice with 20 mL of TBS (20 mM Tris, 150 mM NaCl, pH 7.4), and washed once with 5 mL of FA lysis buffer (50 mM HEPES-KOH, pH 7.4; 150 mM NaCl; 1 mM EDTA, pH 8.0; 1% Triton X-100; 0.1% sodium deoxycholate; 0.1% SDS). Cell pellets were resuspended in 0.5 mL of FA lysis buffer, transferred to a 1.5 mL microtube, and collected by centrifugation at 2,380 × *g* for 1 min at 4°C.

Cells were resuspended in 400 μL of FA lysis buffer containing 1 mM PMSF and 1× Complete EDTA-free solution, transferred to a 2 mL tube (STC-0250, Yasui Kikai), and mixed with 1.5 g of 0.5 mm Zirconia/silica beads. Cells were disrupted using a Multibead Shocker (Yasui Kikai) for 20 cycles of 30 sec on at 2,700 rpm and 30 sec off at 2°C. A burned 23–26-G needle was used to make a few holes at the bottom of the tube, which was inserted into a 5 mL tube and centrifuged at 1,000 × *g* for 30 sec at 4°C. The cell lysate was transferred to a 1.5 mL TPX tube. Chromatin was sheared by sonication at 4°C at high power with ice on a Bioruptor for three sets of 5 cycles of 30 sec on and 30 sec off. Sheared chromatin samples were centrifuged at 20,380 × *g* for 15 min at 4°C and recovered with the same volume of FA lysis buffer. A total of 10 μL was set aside as input DNA.

A total of 320 μL was transferred to a low-binding tube (Costar) and incubated with 10 μL of Dynabeads Protein G (Thermo Fisher) for 1 hr at 4°C, after which the precleared samples were transferred to a new low-binding tube. Then, 2.1 μg of anti-Flag antibody (M2; Sigma‒ Aldrich) was added to each tube, and the mixture was incubated for 3 hrs at 4°C. Chromatin samples were incubated with an antibody and then incubated overnight at 4°C with 12 μL of Dynabeads Protein G, which was preincubated with 5 mg/mL BSA solution in PBS. Beads were washed twice with 0.5 mL of FA lysis buffer, twice with 0.5 mL of FA lysis buffer supplemented with 360 mM NaCl, twice with 0.5 mL of ChIP wash buffer (10 mM Tris-HCl (pH 8.0), 250 mM lithium chloride, 1 mM EDTA (pH 8.0), 0.5% NP-40 and 0.5% sodium deoxycholate), and twice with 0.5 mL of 1× TE buffer. After the buffer was completely removed, 54 μL of ChIP elution buffer (50 mM Tris-HCl (pH 8.0), 10 mM EDTA, and 1% SDS) was added, and the immunoprecipitates were eluted from the beads by incubating for 15 min at 65°C. Fifty μL of eluate was transferred to a new tube and mixed with 150 μL of TE containing 1% SDS. Ten μL of input sample was mixed with 190 μL of TE containing 1%. To each sample, RNaseA was added at a final concentration of 50 μg/mL. These samples were incubated overnight at 65°C for reverse crosslinking and RNA digestion. Then, proteinase K was added at a final concentration of 0.75 mg/mL, and the mixture was incubated for 2 hrs at 50°C. DNA was purified using a QIAquick PCR Purification Kit (QIAGEN). DNA was eluted with 100 μL of dH_2_O.

qPCR was performed on a Thermal Cycler Dice Real Time System II (TaKaRa) with THUNDERBIRD SYBR qPCR Mix or Next SYBR qPCR Mix (TOYOBO) according to the manufacturers’ instructions. One of the input samples was diluted serially to 1/20, 1/200, 1/2,000 and 1/20,000 dilutions, which were subsequently used to construct the standard curve. All the input samples were subsequently diluted 400-fold. IP samples were diluted 10-fold. qPCR was performed with all the diluted samples as well as the standard. The primers used are as follows: in Fig. 3A (i) 5’-AAAATCTCGACCCTTTGGAAGA and 5’- AATGAACCATCGCCAGCAC; Fig. 3A (ii) 5’-GTGCAAAAGACAAATGGATGG and 5’- GGTGCCGTAAATGCAAAACA; Fig. 3A (iii) 5’-TGCAAAGATGGGTTGAAAGAGA and 5’-TTTGCCGCTCTGATGGTG; Fig. 3A (iv) 5’-AACGCGGTGATTTCTTTGCT and 5’- TTGCTAGCCTGCTATGGTTCA; Fig. 3B 5’-ACGAACTTAAGGCCGCAGTA and 5’- CTTCGACAGGTTCCATAACG; Fig. 3C 5’-GCTGGGTCTCACGATCCATATCAG and 5’- TCTTGTTTCGGTGAGTTGGACAGATC. The amount of DNA in each sample was subsequently determined relative to the standard. After considering dilution factors, % input was expressed for each time point of each culture as the amount of DNA in the IP relative to that in the input.

### Fluorescence-activated cell sorter analysis

Approximately 1 × 10^7^ cells were collected at different time points during time course experiments. Cells were fixed with 200 μL of 70% ethanol at −20°C overnight. Cells were collected, resuspended in 200 μL of 50 mM sodium citrate at pH 7.5 containing 0.25 mg/ml RNaseA (Macherey-Nagel) and incubated for 1 hr at 50°C. Then, 100 μL of 50 mM sodium citrate pH 7.5 containing 1 mg/ml Proteinase K (Nacalai) was added and cells were incubated for 1 hr at 50°C. 300 μL of 50 mM sodium citrate pH 7.5 containing 4 μg/ml propidium iodide (Sigma) was added. Cells were sonicated and diluted with 50 mM sodium citrate pH 7.5 containing 4 μg/ml propidium iodide when necessary and were analyzed using a BD Accuri C6 Plus Flow Cytometer (BD Bioscience).

### Quantification and statistical analysis

Statistical analyses were performed using GraphPad Prism, as described in Figure legends.

## RESULTS

### CAF-1 suppresses HR-mediated rDNA instability in response to Fob1-dependent replication fork arrest

To test whether Cac1, Cac2 and Cac3 function as a complex in maintaining rDNA stability, we compared the degree of rDNA instability in *cac1*Δ, *cac2*Δ, and *cac3*Δ single mutants. We isolated genomic DNA from these mutants, separated it by pulsed-field gel electrophoresis (PFGE), and examined the size heterogeneity of chr XII carrying the rDNA array. The wild-type (WT) strain showed homogeneous chr XII bands, while all single *caf-1* mutants showed extremely smeared bands to the degree seen in the *sir2*Δ mutant (Fig. 1B). Consistent with our previous finding (50), the absence of the subunits of budding yeast CAF-1 induced both contraction and expansion of the chromosomal rDNA cluster.

Cells lacking any subunit of the budding yeast CAF-1 showed increased ERC formation (Fig. 1C, 1D and Supplementary Fig. S1A). ERC levels in *cac1*Δ and *cac2*Δ single mutants were ∼3-fold greater than those in the WT strain and their levels were comparable to that in the *sir2*Δ mutant (Fig. 1C, 1D). The *cac3*Δ mutant showed less severe rDNA instability than *cac1*Δ and *cac2*Δ mutants did but still exhibited a ∼1.7-fold increase in the ERC level, compared to the WT strain (Fig. 1C, 1D). While monomeric ERCs were predominant in the WT, the *caf-1* mutants produced more dimeric and multimeric ERCs (Fig. 1E). In both PFGE and ERC analyses, the phenotypes of double and triple *caf-1* mutants were similar to those of *cac1*Δ and *cac2*Δ single mutants (Fig. 1C–1E). Therefore, Cac1, Cac2 and Cac3 function as a complex to stabilize the rDNA by suppressing chromosomal rDNA copy number changes and the production of ERCs, particularly multimeric ERCs.

To elucidate how CAF-1 maintains rDNA stability, we examined whether rDNA instability in the *cac1*Δ mutant depends on Fob1 and the HR protein Rad52. We constructed a diploid strain heterozygous for *cac1*Δ, *fob1* and *rad52*Δ, isolated haploid clones by tetrad dissection, and analyzed their rDNA stability by PFGE. The *cac1*Δ mutant showed smeared chr XII bands, but this smearing was suppressed by an introduction of a *fob1* mutation, suggesting that rDNA instability in the *cac1*Δ mutant depends on Fob1 (Fig. 1F), consistent with a previous finding (50). Smearing of the chr XII bands in the *cac1*Δ mutant was also suppressed by *rad52*Δ (Fig. 1F). Furthermore, ERC production in the *cac1*Δ mutant was suppressed by both *fob1* and by *rad52*Δ (Fig. 1G, 1H). Similar suppression of rDNA instability by *fob1* and *rad52*Δ was observed in *cac2*Δ and *cac3*Δ mutants (Supplementary Fig. S1B–S1E). Thus, CAF-1 suppresses Rad52-mediated rDNA instability in response to Fob1-dependent replication fork arrest at the RFB.

### CAF-1 stabilizes the rDNA in a manner both dependent and independent of repression of transcription from E-pro

CAF-1 is involved in gene silencing at telomeres and silent mating type loci, as well as in silencing marker genes inserted in rDNA (33,51,52). We next examined whether CAF-1 is involved in the regulation of transcription from the native noncoding RNA promoter E-pro. We extracted total RNA from WT, *cac1*Δ, *cac2*Δ and *cac3*Δ mutants, separated RNA by agarose gel electrophoresis under denaturing conditions, and detected transcripts from E-pro by Northern blotting with strand-specific probes (Fig. 2A). The *sir2*Δ mutant, in which transcription from E-pro is derepressed (24), was analyzed in parallel.

**Figure 2.**
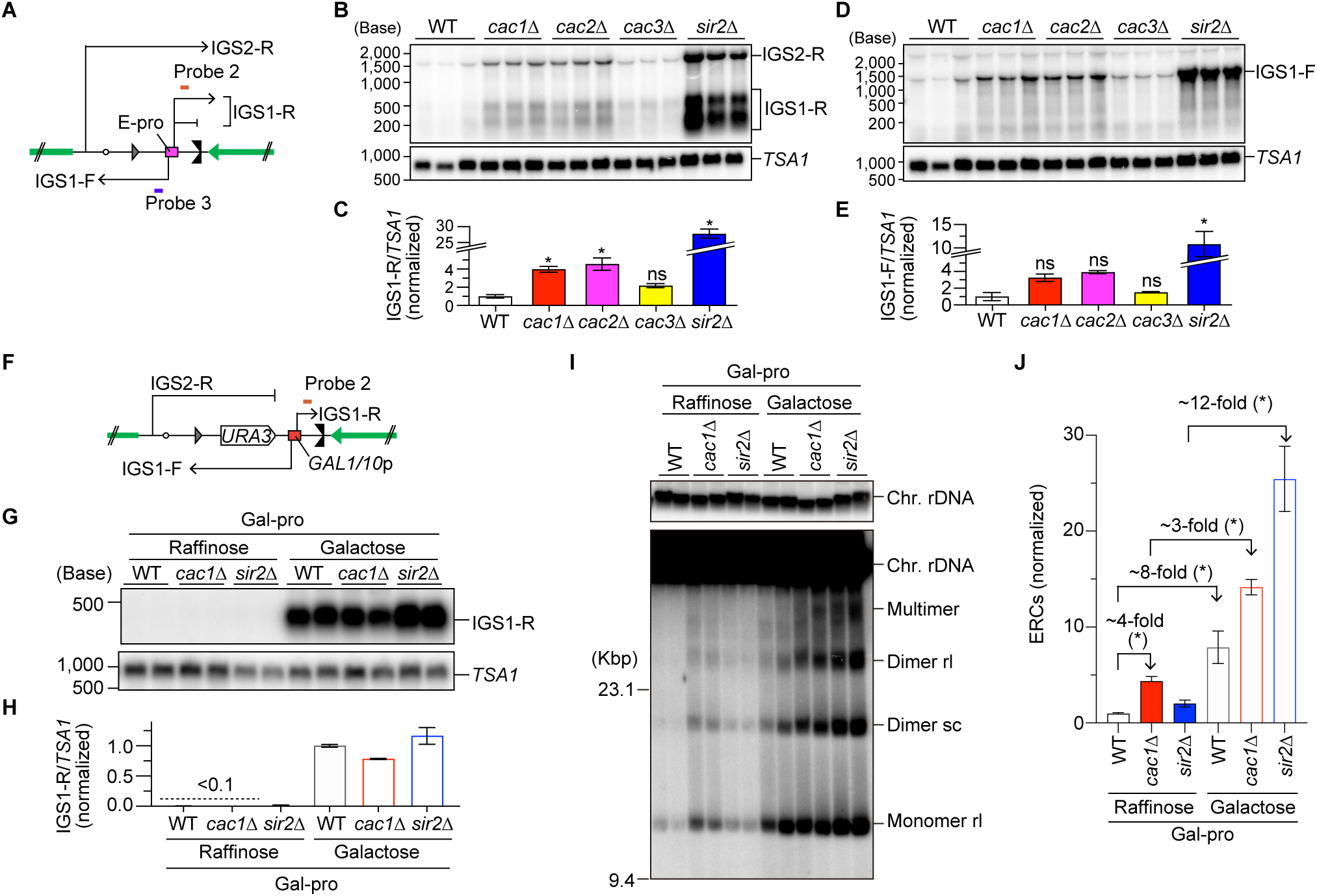
CAF-1 stabilizes the rDNA through both E-pro transcription-dependent and - independent mechanisms. **(A, F)** Positions of probes 2 and 3 used for the detection of noncoding RNA by Northern blotting in strains carrying the native E-pro (A) or Gal-pro (F). **(B, D, G)** Northern blot analysis of noncoding RNAs transcribed from the intergenic regions in the rDNA. Total RNA was isolated from three independent clones of strains carrying E-pro (B, D) and two independent clones of Gal-pro strains grown in the presence of either raffinose or galactose in the media (G). RNA was separated by size on formaldehyde-agarose gels, followed by Northern blotting. In (B) and (G), IGS1-R transcripts were detected with probe 2. In (D), IGS1-F transcripts were detected with probe 3. The membranes were reprobed for *TSA1* transcripts as a control for loading. **(C, E, H)** Quantitation of noncoding RNA. In (C) and (E), signals corresponding to IGS1-R transcripts in (B) and IGS1-F transcripts in (D) were quantified relative to *TSA1*, respectively, which were normalized to the average of WT clones. In (H), IGS1-F transcripts were normalized to *TSA1* and expressed relative to the average of Gal-pro WT clones grown in the presence of galactose. Bars show the mean ± s.e.m. (C, E) and the mean and the range from two independent experiments (H). Transcript levels were compared by one-way ANOVA, followed by Tukey’s multiple comparisons test (C, E); *, statistically significant difference between WT and mutants (p < 0.05); ns, no significant difference (p > 0.05). **(I)** ERC detection. DNA was isolated from independent clones of the indicated strains in the Gal-pro background and separated by agarose gel electrophoresis, followed by Southern blotting with the probe 1, as shown in Fig. 1A. Chromosomal rDNA and different forms of ERCs are indicated. rl and sc indicate relaxed and supercoiled ERCs, respectively. Supercoiled monomers ran off from the gel in the electrophoresis condition under these conditions. The sizes of lambda Hind III DNA markers are indicated. The top panel shows a short exposure of chromosomal rDNA signals. **(J)** Quantitation of ERCs. ERCs were quantified from three independent clones of the indicated strains, including two clones in (H) and a third clone analyzed on a separate gel. ERC levels were compared by the two-sided Student’s t-tests. Bars show the mean ± s.e.m. Asterisks above brackets indicate statistically significant differences between strains that were grown in the indicated growth conditions (*p* < 0.05).

IGS1-R transcripts of ∼300–650 nt in size were detected in the *sir2*Δ mutant, consistent with previous findings (53,54), and their level in the *sir2*Δ mutant was ∼30-fold greater than that in WT cells (Fig. 2B, 2C). The level of IGS1-R transcripts was greater in all *caf-1*Δ mutants than in WT cells, but not as high as those in the *sir2*Δ mutant (Fig. 2C). IGS1-F transcripts of the size ranging from ∼800 to ∼1,200 nt were detected in the *sir2*Δ mutant at a level >10-fold higher than that in WT cells (Fig. 2D, 2E), consistent with previous findings (53,54). Compared to WT, the level of IGS1-F transcripts in *caf-1* mutants was higher, but the increase was less pronounced than that in the *sir2*Δ mutant (Fig. 2D, 2E). For both the IGS1-R and IGS1-F transcripts, *cac1*Δ and *cac2*Δ mutants showed stronger phenotypes than did the *cac3*Δ mutant (Fig. 2B–2E). These results indicate that CAF-1 functions to repress transcription from E-pro, but its contribution is smaller than that of Sir2.

To test the causal relationship between transcription from E-pro and rDNA instability in the *caf-1* mutant, we used the strain that was engineered to have a galactose-inducible, bi-directional *GAL1/10* promoter in place of E-pro in all the rDNA copies, which is hereafter referred to as the Gal-pro strain (Fig. 2F) (24). In Gal-pro WT, *cac1*Δ, and *sir2*Δ strains, transcription from the *GAL1/10* promoter was maintained at an extremely low level when cells were grown in the presence of raffinose but was induced when galactose was added to the media (Fig. 2G, 2H). ERC production was strongly suppressed in both the WT and *sir2*Δ mutant strains in the Gal-pro background when grown in the presence of raffinose (Fig. 2I, 2J). In contrast, in the Gal-pro *cac1*Δ mutant, the level of ERCs was ∼4-fold higher than that in the Gal-pro WT strain in the presence of raffinose, where IGS1-R transcripts were not detected (Fig. 2G–2J, Gal-pro *cac1*Δ in raffinose vs. Gal-pro WT in raffinose). Thus, defects that were independent of enhanced transcription near the RFB contributes to ERC production in the *cac1*Δ mutant.

The addition of galactose to the media induced ERC production by 8- and 12-fold in the Gal-pro WT and *sir2*Δ mutant strains, respectively, compared to the same strains when grown in the presence of raffinose (Fig. 2I, 2J). This indicates that ERC production in WT and *sir2*Δ mutant predominantly depends on the transcriptional activity near the RFB, consistent with previous findings (24,55). In the Gal-pro *cac1*Δ mutant, the addition of galactose to the media induced ERC production by ∼3-fold, compared to that when grown in the raffinose-containing media, but the degree of its induction was lower than that in the Gal-pro WT and *sir2*Δ cells (Fig. 2I, 2J). Taken together, these findings suggest that CAF-1 stabilizes the rDNA in a manner that is both dependent and independent through its function to repress transcription at the E-pro locus.

### CAF-1 suppresses end resection of replication-coupled DSBs in the rDNA

To test whether enhanced E-pro activity resulted in cohesin dissociation from the CAR, we conducted chromatin immunoprecipitation and quantitative PCR in WT, *cac1*Δ and *sir2*Δ strains expressing the cohesin subunit Mcd1 tagged with a Flag epitope tag (Fig. 3A). Cohesin was associated with the positive control region near a centromere on chr VI (Fig. 3B) (56), but not with the negative control region near *YEL008W* on chr V (Fig. 3C), where cohesin association was not observed in a previous chromatin immunoprecipitation sequencing study (57). In the rDNA region, cohesin was most enriched in the CAR in exponentially growing WT cells, and cohesin association was decreased in the *sir2*Δ mutant (Fig. 3A, ii), consistent with previous findings (10). Although the *cac1*Δ mutant showed enhanced transcription from E-pro, compared to that in WT cells (Fig. 2B–2E), it did not result in the inhibition of cohesin association with the CAR (Fig. 3A).

**Figure 3.**
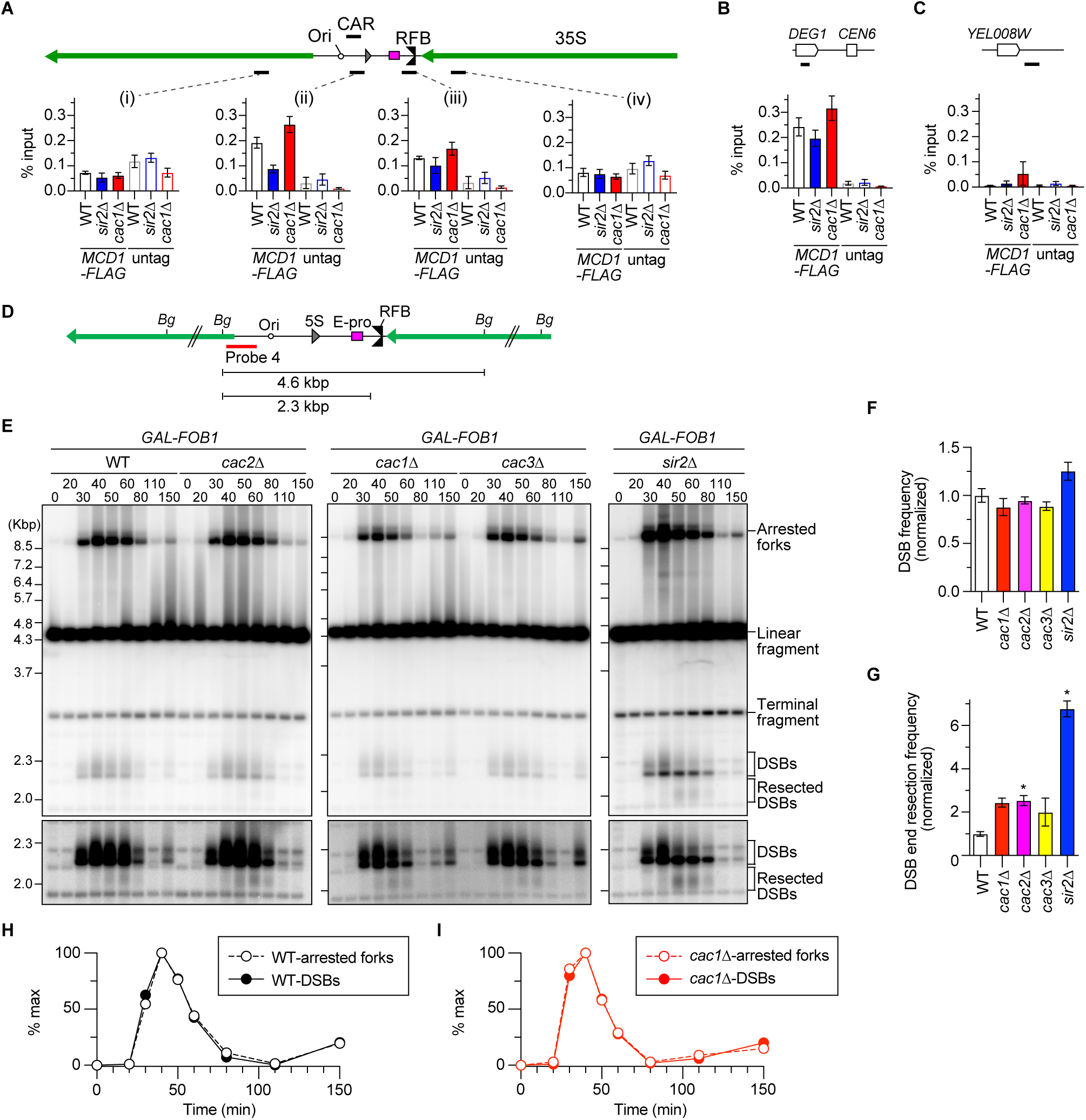
CAF-1 suppresses DSB end resection. **(A, B, C)** ChIP‒qPCR analyses of Mcd1-Flag in the indicated strains. The regions where cohesin enrichments were examined are indicated as black bars. Four different regions in the rDNA region (i)–(iv) in (A), the positive control region near *CEN6* in (B) and the negative control region near *YEL088W* on chr V (C) were examined. Untagged strains were used as controls. ChIP was performed using an antibody against a Flag epitope tag. The enrichment of chromatin in the indicated regions is expressed as the % input (IP/whole-cell extract). The bars show the mean ± s.e.m. from three independent cultures of each strain. **(D)** Restriction map and the positions of the probe 4 used for DSB analyses. The restriction sites for Bgl II (Bg) are indicated. **(E)** DSB assay. Time-course experiments were conducted by arresting the indicated strains in the *GAL-FOB1* background in G1 with alpha factor, releasing cells into fresh media and collecting cells at different time points, as described in Supplementary Fig. S2A. Genomic DNA was isolated, digested with Bgl II, separated by size, and analyzed by Southern blotting with probe 4, as shown in (D). Arrested forks, linear fragments, DSBs and resected DSBs are indicated. Terminal fragments indicate the telomere-proximal rDNA repeat. Long exposure panels around DSBs and resected DSBs are shown below. **(F)** Quantitation of DSBs. The frequency of DSBs was determined by quantifying the maximum signal of DSBs relative to the maximum signal of arrested forks in each time course. The average DSB frequency from three independent time course experiments was determined and normalized to the average of WT clones (bars show the mean ± s.e.m.). Multiple comparisons were performed by one-way ANOVA, followed by Tukey’s multiple comparisons test. The difference in DSB frequencies between WT and all other mutants was not statistically significant. **(G)** Quantitation of resected DSBs. The frequency of resected DSB was determined by quantifying the maximum signal of resected DSBs relative to the maximum signal of DSBs in each time course. The average frequency of resected DSBs from three independent time course experiments was determined and normalized to the average of WT clones (bars show the mean ± s.e.m.). Multiple comparisons were performed by one-way ANOVA, followed by Tukey’s multiple comparisons test; *, statistically significant difference (p < 0.05). **(H, I)** Timing of appearance and disappearance of arrested forks and DSBs. Arrested forks and DSBs are expressed as percent of maximum values in one of the three independent time course experiments of *GAL-FOB1* WT (H) and *cac1Δ* strains (I).

We next examined whether CAF-1 impacts DSB formation and its repair by conducting time course experiments during synchronous S phase progression in cells expressing the *FOB1* gene under the control of the *GAL1* promoter (Fig. S2A). We prepared genomic DNA, digested it with Bgl II (Fig. 3D), and separated it by conventional agarose gel electrophoresis, followed by detection of arrested forks, DSBs, and resected DSBs by Southern blotting (Fig. 3E). The DSB frequency was estimated by normalizing the maximum level of DSBs relative to that of arrested forks in each time course. CAF-1 deficiency did not significantly impact DSB levels (Fig. 3F).

The frequency of DSB end resection was determined by normalizing the maximum level of resected DSB signal to that of DSBs in each time course. Consistent with our previous finding (23), the absence of Sir2 led to a ∼7-fold increase in the frequency of DSB end resection (Fig. 3G). Cells lacking any subunits of CAF-1 showed ∼2–3-fold increase in the frequency of DSB end resection, compared to that in WT cells, but this increase was not as extensive as that seen in cells lacking Sir2 (Fig. 3G). Because *GAL-FOB1* WT and *caf-1* mutants showed comparable synchrony and timing in their progression through S phase (Supplementary Fig. S2B), enhanced DSB end resection in *GAL-FOB1 caf-1* mutants was unlikely caused by differences in the efficiency of release from G1 arrest by alpha factor. Taken together, these findings suggest that CAF-1 suppresses end resection of DSBs formed at the RFB.

### The MRX complex is important for DSB repair and ERC production in the *caf-1* mutant

The MRX complex, along with its cofactor Sae2, initiates short-range end resection of double-ended DSBs introduced by endonucleases, and Exo and Sgs1-Dna2 carry out long-range resection (15,58,59). When *MRE11* or *SAE2* was deleted from *GAL-FOB1 cac1*Δ cells, resected DSBs were barely detectable at the RFB (Fig. 4A and 4D, Supplementary Fig. S3A and S3D). Therefore, the MRX complex and Sae2 are responsible for initiating end resection of replication-coupled DSBs at the RFB in the *cac1*Δ mutant.

**Figure 4.**
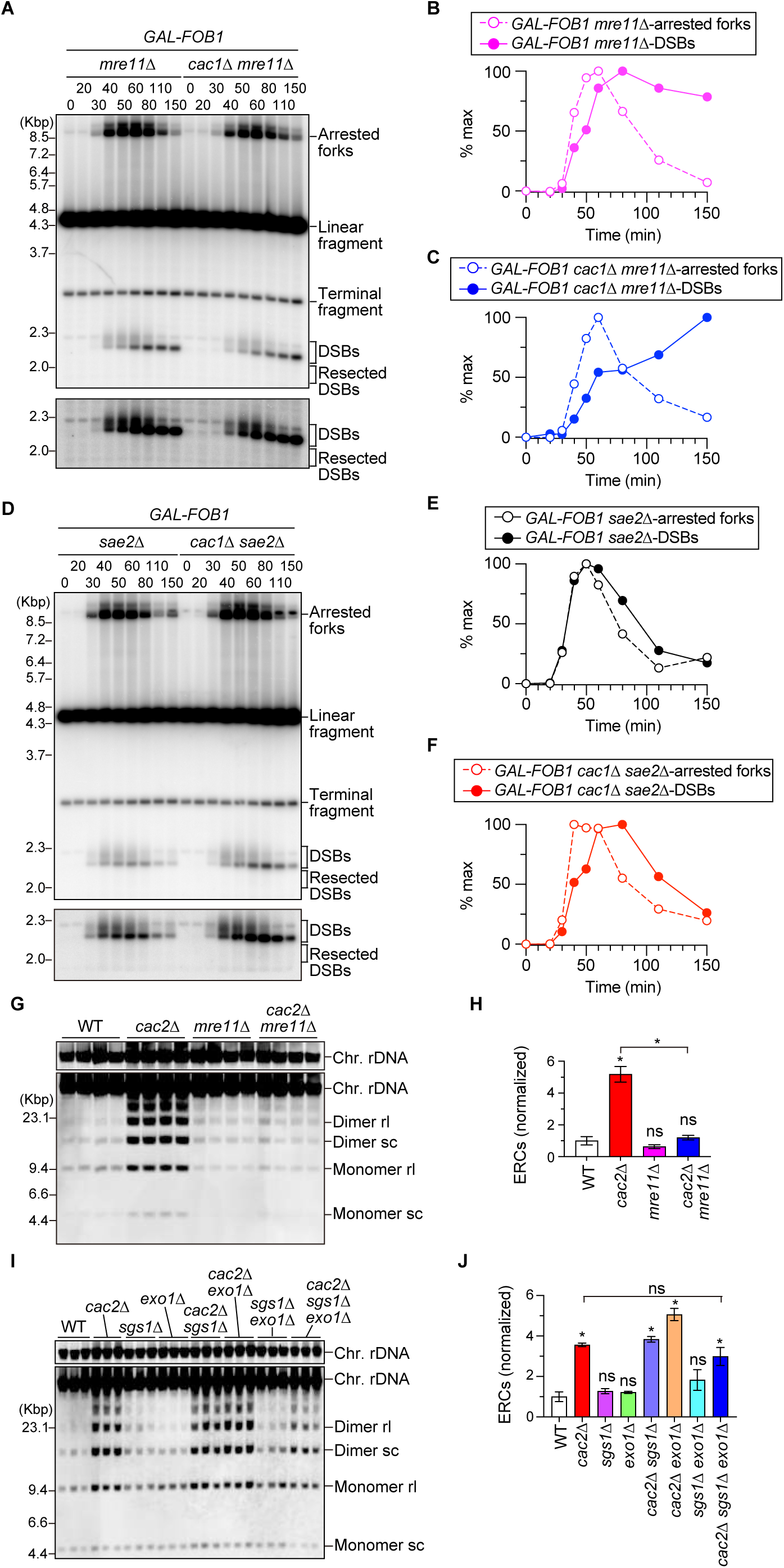
The MRX complex is required for DSB repair and ERC production in the *caf-1* mutant. **(A, D)** DSB assay. Time-course experiments were conducted by arresting the indicated strains in the *GAL-FOB1* background in G1 with alpha factor, releasing cells into fresh media and collecting cells at different time points, as described in Supplementary Fig. S2A. Genomic DNA was isolated, digested with Bgl II, separated by size, and analyzed by Southern blotting with probe 4, as shown in Fig. 3D. Arrested forks, linear fragments, DSBs, and resected DSBs are indicated. Terminal fragments indicate the telomere-proximal rDNA repeat. Long exposure panels around DSBs and resected DSBs are shown below. **(B, C, E, F)** Timing of appearance and disappearance of arrested forks and DSBs. Arrested forks and DSBs are expressed as percent of maximum values in one of the two independent time course experiments of the indicated strains. **(G, I)** ERC detection. DNA was isolated from four independent clones (G) and three independent clones (J) of the indicated strains and separated by agarose gel electrophoresis, followed by Southern blotting with the probe 1, as shown in Fig. 1A. Chromosomal rDNA and different forms of ERCs are indicated. rl and sc indicate relaxed and supercoiled ERCs, respectively. The sizes of lambda Hind III DNA markers are indicated. The top panel shows a short exposure of chromosomal rDNA signals. **(H, J)** Quantitation of ERCs. In (H) and (J), ERCs were quantified from (G) and (I), respectively. ERC levels were compared between WT and mutants by one-way ANOVA, followed by Tukey’s multiple comparisons test. Bars show the mean ± s.e.m. Asterisks indicate statistically significant differences between WT and mutant strains (p < 0.05). ns indicates no significant difference (p > 0.05).

In *GAL-FOB1* WT cells, DSB signal appeared and disappeared at the timing similar to that of arrested fork signal (Fig. 3E, 3H). *GAL-FOB1 cac1*Δ cells also showed efficient DSB repair, as the timing of disappearance of DSB signal was similar to that of arrested fork signal (Fig. 3I). On the contrary, in *GAL-FOB1* cells lacking Mre11, DSBs accumulated at the end of time courses when most cells had completed bulk DNA synthesis and arrested forks had almost disappeared (Fig. 4A, 4B and Supplementary Fig. S3A, S3B), consistent with a previous finding (17,23). *GAL-FOB1 cac1*Δ *mre11*Δ cells also showed accumulation of DSBs at the end of time courses (Fig. 4A, 4C and Supplementary Fig. S3A, S3C). When *SAE2* was deleted from *GAL-FOB1* WT and *cac1*Δ cells, there was a slight delay in the disappearance of DSB signals, compared to that of arrested fork signals, but DSB signals gradually decreased by the end of time course, unlike DSBs that remained accumulated in cells lacking Mre11 (Fig. 4A–4F and Supplementary Fig. S3). These findings demonstrate that the MRX complex is essential for DSB repair but its function to carry out DSB end resection may be less important for DSB repair.

Enhanced ERC production in the *cac2*Δ mutant was suppressed by deletion of *MRE11* to the level seen in the *mre11*Δ single mutant, which was comparable to or slightly lower than that in WT (Fig. 4G, 4H). Exo1 and Sgs1-Dna2 function redundantly to carry out long-range DSB end resection (15,60). ERC levels in the *cac2*Δ mutant appeared slightly lowered by deletion of *SGS1* and *EXO1*, but this difference was not statistically significant (Fig. 4I, 4J). Taken together, these findings suggest that the MRX complex is essential for DSB repair and prevention of rDNA instability by limiting ERC production.

### The interaction of CAF-1 with PCNA is critical for rDNA stabilization

CAF-1 is known to be recruited through its interaction with PCNA and deposit histones onto the newly synthesized DNA (31–33). To test whether this recruitment is important for CAF-1’s function in rDNA, we examined the effect of mutations that compromise the interaction between CAF-1 and PCNA. We introduced an empty plasmid or plasmids carrying the *CAC1* or *cac1-20* allele that had a mutation in the PCNA-binding motif of the Cac1 protein (61) and analyzed rDNA stability by PFGE and ERC analyses.

The *cac1*Δ mutant transformed with an empty vector showed smeared chr XII bands, while the *cac1*Δ mutant complemented with a plasmid carrying *CAC1* showed sharp bands (Fig. 5A). Expression of *cac1-20* in the *cac1*Δ mutant resulted in chr XII bands that appeared less smeared than those in the *cac1*Δ mutant with an empty vector, although the bands were still broader than those in the *cac1*Δ mutant expressing *CAC1* (Fig. 5A). Expression of *CAC1* led to lowering the level of ERCs in the *cac1*Δ mutant carrying an empty plasmid. However, expression of *cac1-20* in the *cac1Δ* mutant did not cause a reduction in the ERC level, compared to that in the *cac1*Δ mutant expressing *CAC1* (Fig. 5B, 5C). Therefore, the *cac1-20* mutant defective in its binding to PCNA displayed rDNA instability. As the level of Cac1-20 was comparable to that of Cac1 (Fig. 5D, 5E), enhanced rDNA instability in the *cac1-20* mutants was unlikely caused by reduced expression due to its mutation.

**Figure 5.**
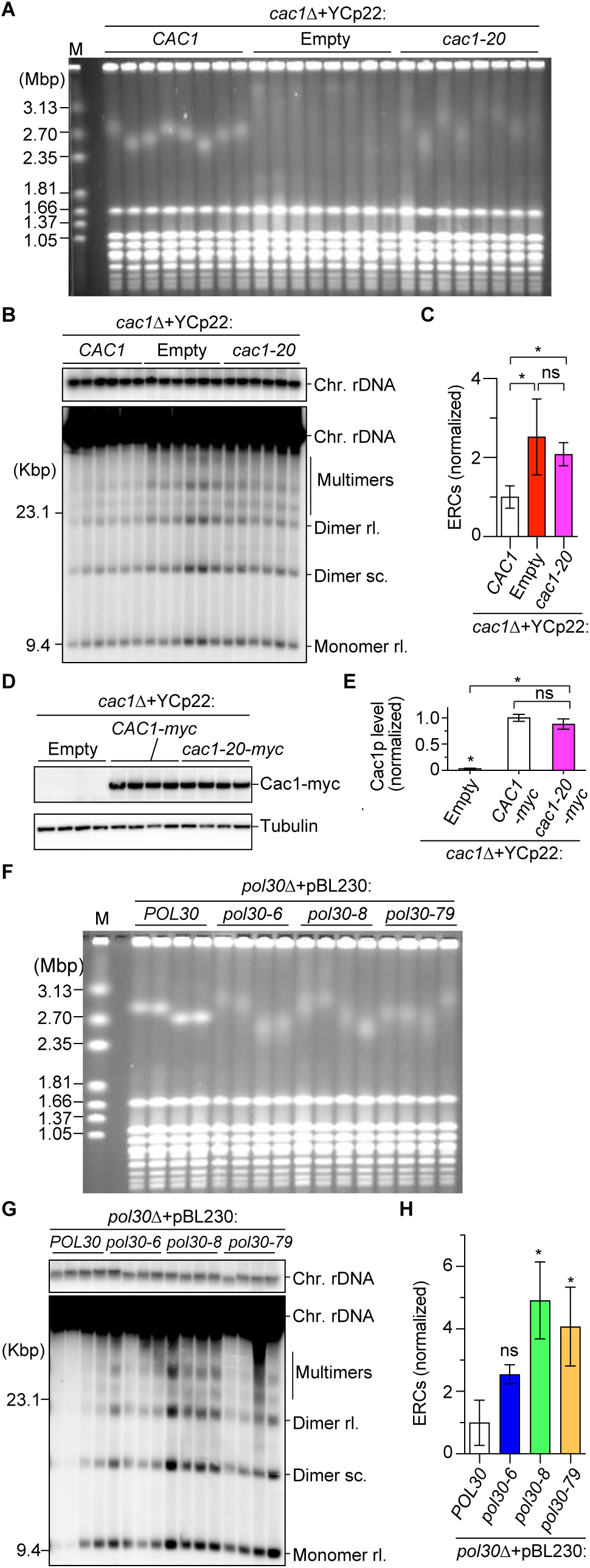
Interaction with PCNA is important for CAF-1 to stabilize the rDNA. **(A, F)** PFGE analysis to examine the size heterogeneity of chr XII. DNA from independent clones of the indicated strains was separated by PFGE and stained with EtBr. M indicates *H. wingei* chromosomal DNA markers. **(B, G)** ERC detection. DNA was isolated from 6 independent clones (B) and four independent clones (G) of the indicated strains and separated by agarose gel electrophoresis, followed by Southern blotting with the probe 1, as shown in Fig. 1A. Genomic rDNA and different forms of ERCs are indicated. M indicates lambda Hind III DNA markers. The top panel shows a short exposure of genomic rDNA signals. **(C, H)** Quantitation of ERC levels. ERCs in (B, G) were quantified and their levels were normalized to those of the *cac1*Δ strain complemented with a plasmid carrying *CAC1* in (C) and the *pol30*Δ strain complemented with a plasmid carrying *POL30* in (H). ERC levels among different strains were compared by one-way ANOVA, followed by Tukey’s multiple comparisons test. Bars show the mean ± s.e.m. Asterisks above bars and above brackets indicate statistically significant differences between WT and mutant strains and between mutant strains, respectively (*p* < 0.05). ns indicates no significant difference (p > 0.05). **(D)** Western blot of Cac1 that is C-terminally tagged with a nine Myc epitope and tubulin from four independent clones of the *cac1*Δ strains that were transformed with a YCp22 vector or the vectors expressing either Cac1-9myc or Cac1-20-9myc. **(E)** Quantitation of Cac1-myc protein levels. The levels of Cac1-9myc and Cac1-20-9myc were quantified in the indicated strains relative to tubulin, which was normalized to the average of the *cac1*Δ clones expressing Cac1-9myc. Bars show the mean ± s.e.m. Multiple comparisons were performed by one-way ANOVA, followed by Tukey’s multiple comparisons test; *, statistically significant difference (p < 0.05); ns, no significant difference (p > 0.05).

We also analyzed the effects of three mutations in *POL30*, which encodes PCNA: *pol30-6*, *pol30-8*, and *pol30-79*, all of which results in a substantial loss of the ability of PCNA to bind to Cac1p (33). Because the *POL30* gene is essential, we constructed a diploid strain heterozygous for *pol30*Δ, transformed it with plasmids carrying wild-type *POL30* or mutant alleles, and isolated haploid *pol30*Δ mutants inheriting plasmids by tetrad dissection. In PFGE, the *pol30*Δ mutant complemented with *POL30* showed sharp chr XII bands, but the *pol30*Δ mutants expressing *pol30-6*, *pol30-8* or *pol30-79* showed smeared chr XII bands (Fig. 5F). Moreover, compared with the *pol30*Δ mutant complemented with *POL30*, the *pol30*Δ mutants expressing mutant alleles showed ∼2.5-fold or even higher level of ERCs (Fig. 5G, 5H). rDNA instability seen in these mutants was unlikely caused by reduced expression of these mutant proteins, because the previous study demonstrated that the level of Pol30-6, Pol30-8 and Pol30-79 proteins bound to chromatin are comparable to that of Pol30 (33). Taken together, these findings suggest that the interaction of CAF-1 with PCNA, which is necessary for the deposition of histones onto newly synthesized DNA, is important for the maintenance of rDNA stability.

### CAF-1-mediated chromatin assembly facilitates properly spaced lagging strand synthesis in the rDNA

During DNA replication *in vivo*, Okazaki fragment synthesis is interlinked with chromatin assembly (39). Thus, analysis of Okazaki fragments allows us to assess nucleosome assembly *in vivo*. To evaluate Okazaki fragment synthesis in the rDNA region, we used the strains carrying a doxycycline-repressible allele of *CDC9* that encodes a DNA ligase joining Okazaki fragments, isolated genomic DNA from cultures where *CDC9* was either shut off or kept expressed, and separated equal amounts of genomic DNA by agarose gel electrophoresis under denaturing conditions, followed by Southern blotting with an rDNA probe.

Upon *CDC9* inhibition, WT cells showed a ladder of bands that appeared to be sized according to the nucleosomal repeat (165 bp) (Fig. 6A and Supplementary Fig. S4), consistent with a previous finding that analyzed Okazaki fragments generated during bulk DNA synthesis (39). When CAF-1 subunits were absent, overall lane signals were reduced, compared to those seen in WT, even though equal amounts of DNA were loaded (Fig. 6A and Supplementary Fig. S4). Previous studies analyzing bulk DNA demonstrated that the absence of CAF-1 subunits completely abolishes the nucleosome-sized periodicity of Okazaki fragments and increases the average length of Okazaki fragments (39–41). However, in our analysis that analyzed the rDNA-specific effect, we still observed periodic signal peaks in *caf-1* mutants at the reduced intensity, compared to those in WT (Fig. 6A and Supplementary Fig. S4). Furthermore, we did not observe clear evidence of lengthening of Okazaki fragments. Nonetheless, these results indicate that the *caf-1* mutants exhibited partial defects in nucleosome assembly in the rDNA.

**Figure 6.**
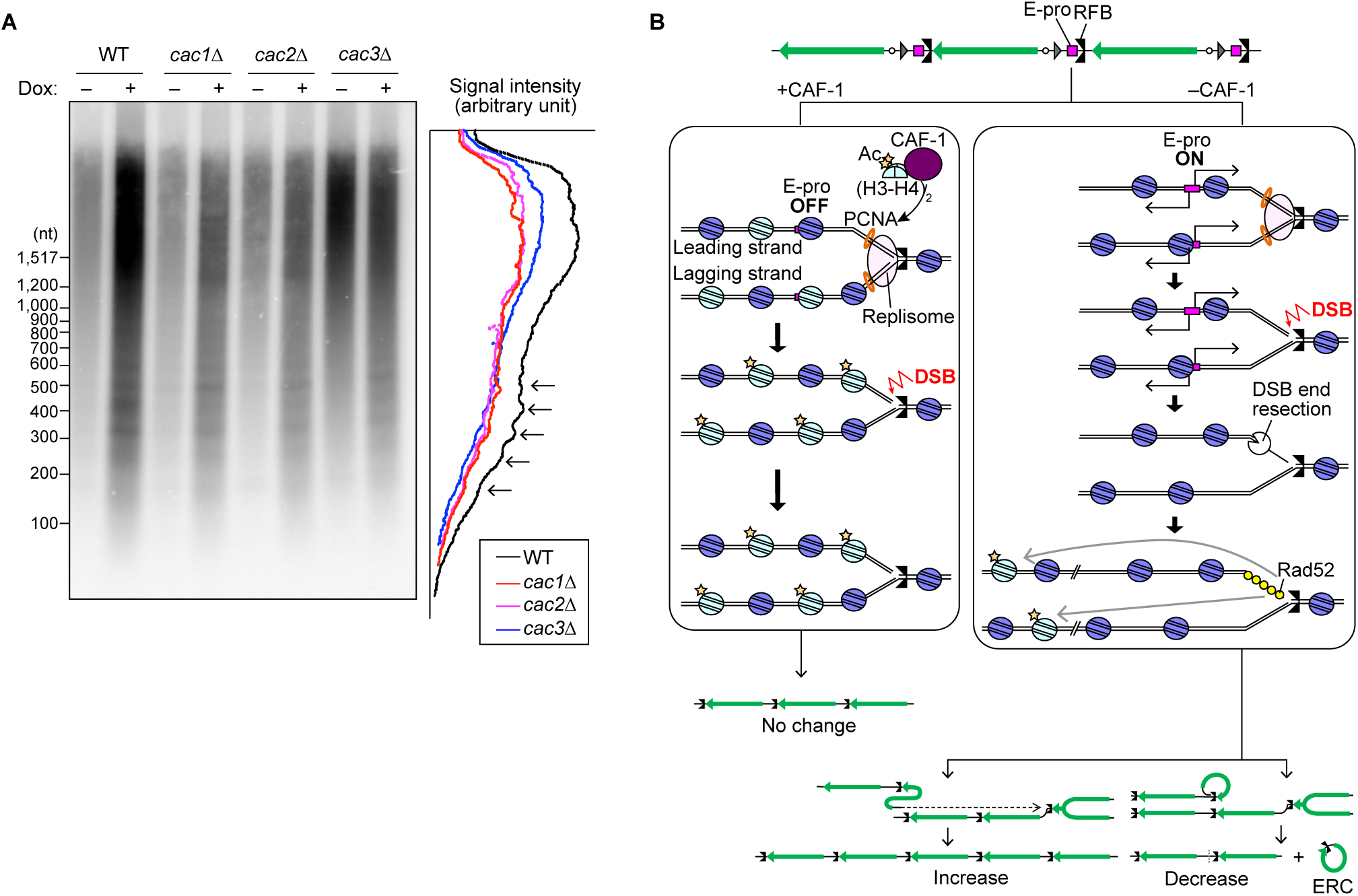
A model for CAF-1-mediated rDNA stabilization. **(A)** The size of Okazaki fragments generated during rDNA replication. WT, *cac1*Δ, *cac2*Δ,and *cac3*Δ cells carrying a doxycycline-repressible allele of *CDC9* were treated or not treated with doxycycline (Dox). Genomic DNA was isolated and separated on a denaturing agarose gel, followed by Southern blotting with a rDNA probe 1, as indicated in Fig. 1A. Lane profiles of the indicated strains, in which *CDC9* was shut off, were included. The positions of the bands of Okazaki fragments are indicated by arrowheads. **(B)** A model for CAF-1-mediated rDNA stabilization. Left: CAF-1 facilitates nucleosome assembly onto replicating DNA by depositing a newly synthesized (H3-H4)_2_ tetramer where H3K56 is acetylated (Ac). The assembly of the properly spaced nucleosome leads to the repression of transcription from E-pro. DSBs formed in this chromatin environment are repaired in a manner that does not involve rDNA copy number changes. Right: The absence of CAF-1 leads to defects in depositing newly synthesized histones. This environment results in the activation of transcription from E-pro, leading to the induction of DSB end resection and HR-mediated DSB repair. The HR protein Rad52 bound to resected DSBs preferentially uses the rDNA copy in a chromatin environment with histone H3K56Ac. Because the density of histone H3 acetylated at K56 is low in the CAF-1-defective strain, Rad52 assembled on the replicated DNA preferentially uses the distant rDNA copy, inducing rDNA instability. This template choice error may not be affected by transcription at the E-pro.

## DISCUSSIONS

Eukaryotic organisms have evolved mechanisms that faithfully maintain genetic and epigenetic information. CAF-1 is one of the factors involved in these processes and ensures the coordination of DNA replication with chromatin assembly (62). In this study, we demonstrated that the absence of CAF-1 caused frequent expansion and contraction of the chromosomal rDNA array as well as enhanced ERC production (Fig. 1). CAF-1 is important for the rDNA stability in *A. thaliana*, but *A. thaliana* lacking CAF-1 leads only to loss of rDNA copies and does not produce ERCs (42–44). The reasons behind these differences remain unclear. rDNA instability in budding yeast *caf-1* mutants depends on Rad52 (Fig. 1F–1H), extending insights from previous studies in *A. thaliana* (43). rDNA instability in budding yeast also depends on Fob1 that is responsible for replication fork arrest at the RFB and subsequent DSB formation (Fig. 1F–1H). Our findings suggest that CAF-1 suppresses HR-mediated chromosomal and extrachromosomal rDNA instability during the repair of DSBs formed in the context of arrested replication forks at the RFB.

CAF-1 is involved in silencing noncoding RNA transcription from the endogenous E-pro (Fig. 2B–2E). Using Gal-pro *caf-1* mutants, we showed that the activation of transcription at the E-pro locus induces ERC production, but these mutants produce ERCs even when this transcription was kept low. Therefore, CAF-1 stabilizes the rDNA in a manner dependent and independent through its function to repress transcription from E-pro (Fig. 2F–2J).

Previous studies demonstrate that the activation of transcription at the E-pro locus prevents cohesin binding to the rDNA (10,23,24). Because a temperature-sensitive mutant of *smc1*, which encodes one of the cohesin subunits, shows frequent loss of a marker gene from the rDNA cluster, a previous model proposes that cohesins bound to the CAR and RFB contribute to restricting the interaction of the DSB end with the misaligned rDNA copy during DSB repair (10). Unexpectedly, the *cac1*Δ mutant, which showed a ∼4-fold increase in the transcription level at the E-pro locus (Fig. 2B–2E), did not show reduced cohesin binding but rather showed a slight increase in cohesin binding to the rDNA (Fig. 3A), indicating that this enhanced transcription is unlikely to affect rDNA instability through regulation of cohesion associations. However, it is known that Eco1-dependent acetylation of a cohesin-subunit Smc3, which is essential for cohesion establishment, is reduced in the *caf-1* mutant (63). Therefore, we cannot exclude the possibility that CAF-1 is involved in the conversion of cohesin bound to the rDNA locus into its establishment mode to stabilize the rDNA.

CAF-1 deficiency led to increased DSB end resection (Fig. 3E, 3G). We previously demonstrated by analyzing the *sir2*Δ mutant that transcription activation at the E-pro locus induces the collision of transcription machineries to the arrested replication forks at the RFB, which recruits Mre11 and induces DSB end resection (23). Therefore, derepression of transcription from E-pro in the *cac1*Δ mutant most likely contribute to enhanced DSB end resection (Fig. 2B–2E, 3F). Alternatively, it is possible that chromatin assembled by CAF-1 at the DSB may prevent DSB end resection by restricting the access by DSB end resection enzymes. The MRX complex was required for DSB end resection, DSB repair, and ERC production in the *caf-1* mutant, but ERC production in the *caf-1* mutant was not affected or only slightly reduced, if any at all, by absence of both Exo1 and Sgs1 (Fig. 4). Thus, long-range resection was not the absolute requirement for ERC production.

CAF-1 prevents Rad52-mediated rDNA instability during repair of replication-coupled DSB repair at the RFB (Fig. 1F–1H). Knockdown of CAF-1 and another histone chaperone ASF1 in human U-2 OS cells leads to reduced HR in response to replication-independent DSBs introduced by an endonuclease I-SceI as well as reduced foci of Rad51 and its loader MMS22L-TONSL upon treatment with DNA damaging agent bleomycin, leading to the model that CAF-1 promotes proper chromatin assembly onto resected DSB ends to promote HR by facilitating Rad51 loading onto resected DSBs (37). The function of CAF-1 in rDNA appears to contrast to that of human CAF-1 in response to replication-independent DSBs, because CAF-1 prevents rDNA recombination (Fig. 1F–1H). Knockdown of ASF1 in human cells additionally shows increased levels of resected DSBs upon bleomycin treatment (37). Thus, human ASF1 is involved in preventing DSB end resection, but it remains to be tested whether CAF-1 also elicit similar function. A recent study in budding yeast has developed a system to introduce replication-coupled DSBs using a Cas9 nickase (21). Absence of Asf1 causes severe growth defects, when replication-dependent DSBs are introduced in the non-rDNA region by Cas9 nickase and guide RNA. CAF-1 deficiency reduces cellular growth but only when combined with another histone chaperone Rtt106 (21). Thus, budding yeast CAF-1 plays a minor role in response to replication-coupled DSBs caused by nicks on the template DNA. These findings raise the possibility that CAF-1 elicits distinct functions in DSB repair, depending on the sources of DSBs.

How does CAF-1 promote rDNA stability independently through its regulation of transcription from E-pro? We analyzed Okazaki fragments during rDNA replication to assess chromatin assembly around the RFB and found that abundance of nucleosome-sized Okazaki fragments decreased in the *cac1*Δ mutant (Fig. 6A and Supplementary Fig. S4), indicating that the nucleosome occupancy behind arrested forks and thus around DSBs at the RFB decreases when CAF-1 is absent (Fig. 6B). CAF-1-mediated assembly of nucleosomes on newly replicated DNA may be important for restricting DSB end resection and preventing the repair of DSBs with the misaligned copy, which needs to be examined in future studies.

CAF-1 loads newly synthesized histones H3-H4, where histone H3K56 is acetylated by the histone chaperone Rtt109 (28–30). Defective acetylation of histone H3K56 renders cells sensitive to DNA damaging agents, suggesting that chromatin with histone H3K56 acetylation facilitates DNA repair (29). We previously demonstrated that the absence of Rtt109, which is responsible for the acetylation of histone H3K56, induces rDNA instability (35). We speculate that during DSB repair, Rad52 preferentially uses a homologous rDNA copy that contains chromatin with acetylated histone H3K56 (Fig. 6B). Deficiency of CAF-1 results in defects in the deposition of nucleosomes with acetylated histone H3K56 on replicating DNA (Fig. 6B, right). Defective nucleosome assembly may derepress transcription from E-pro, which contributes to enhanced DNA end resection and Rad52-mediated repair. The likelihood of having acetylated histone H3K56 is low behind the arrested forks at the RFB. Thus, in cells lacking CAF-1, Rad52 is unlikely to use the rDNA copy at an aligned position for DSB repair due to the absence of H3K56 acetylation but instead preferentially uses a copy that is distant from the DSB site, leading to chromosomal rDNA copy number changes and ERC production (Fig. 6B). Defects in proper assembly of chromatin with H3K56 acetylation may promote frequent usage of the misaligned copy during DSB repair, independently of the transcription status at the E-pro.

## Data availability

All data are available upon request.

## Acknowledgements

We are grateful to Peter Burgers for the plasmids pBL230-0, 6, 8 and 79 and Iestyn Whitehouse for the yIW310 strain. We thank Etsuko Imatani and Akemi Mizuguchi for technical assistances and members of the Kobayashi and Sasaki laboratories for discussions.

## Author contributions

T.K. and M.S. conceived the research. H.F., M.S. and T.K. designed the experiments and analyzed the data. H.F., T.Y., Y.K., N.A., and M.S. performed the experiments. M.S. wrote the manuscript with input from T.K.

## Funding

Japan Society for the Promotion of Science [20H05382 and 24K09417 to M.S. and 17H01443 and 21H04761 to T.K.]; Japan Science and Technology Agency Fusion Oriented REsearch for disruptive Science and Technology, FOREST [JPMJFR214P to M.S.]; Naito Foundation (to M.S.); Japan Agency for Medical Research and Development [JP21gm1110010 to T.K.]; Core Research for Evolutional Science and Technology [JPMJCR19S3 to T.K.], Strategic Research Projects grant from Research Organization of Information and Systems.

## Conflict of interest statement

None declared.

---

**Supplementary Figure 1. ERC detection in the *caf-1* mutants.**

**(A, B, D)** ERC detection. DNA was isolated from the two independent clones (A) and four independent clones (B, D) of the indicated strains and separated by agarose gel electrophoresis, followed by Southern blotting with the probe 1, as shown in Fig. 1A. Chromosomal rDNA and different forms of ERCs are indicated. rl and sc indicate relaxed and supercoiled ERCs, respectively. Supercoiled monomers ran off from the gel in the electrophoresis condition under these conditions. The sizes of lambda Hind III DNA markers are indicated. The top panel shows a short exposure of chromosomal rDNA signals. The DNA sample that was indicated as WT-1 was the same DNA sample isolated from the clone that was loaded a separate gel: WT-1 in (A) is the first WT clone in Fig. 1C; WT-1 in (B) is the first WT clone on the left gel; WT-1 in (D) is the first WT clone on the left gel. These DNA samples were used to normalize ERC levels in the mutants that were analyzed on separate gels.

**(C, E)** Quantitation of ERCs. In (C) and (E), ERCs were quantified from (B) and (D), respectively. Multiple comparisons were performed by one-way ANOVA, followed by Tukey’s multiple comparisons test; *, statistically significant difference (p < 0.05).

**Supplementary Fig. 2. Time course experiments to analyze formation and repair of DSBs.**

(A) Outline of time course experiments.

(B) Cell cycle analysis of two representative time course experiments using *GAL-FOB1* WT, *cac1*Δ, *cac2*Δ, *cac3*Δ, and *sir2*Δ strains by flow cytometry.

**Supplementary Figure 3. Time course experiments to analyze DSB repair in *cac1*Δ *mre11*Δ and *cac1*Δ *sae2*Δ.**

**(A, D)** DSB assay. Time-course experiments were conducted, as described in Supplementary Fig. S2A. Genomic DNA was isolated, digested with Bgl II, separated by size, and analyzed by Southern blotting with probe 4, as shown in Fig. 3D. Arrested forks, linear fragments, DSBs and resected DSBs are indicated. Terminal fragments indicate the telomere-proximal rDNA repeat. Long exposure panels around DSBs and resected DSBs are shown below.

**(B, C, E, F)** Arrested forks and DSBs are expressed as percent of maximum values in one of the two independent time course experiments of the indicated strains.

**Supplementary Figure 4. The size of Okazaki fragments in the *caf-1* mutants.**

Okazaki fragments were analyzed in WT, *cac1*Δ, *cac2*Δ, and *cac3*Δ cells carrying a doxycycline-repressible allele of *CDC9* that were treated or not treated with doxycycline (Dox). Genomic DNA was isolated and separated on a denaturing agarose gel, followed by Southern blotting with probe 1, as indicated in Fig. 1A. Lane profiles of the indicated strains, in which *CDC9* was shut off, were included. The positions of the bands of Okazaki fragments are indicated by arrowheads.

## Notes

### Competing Interest Statement

The authors have declared no competing interest.

### Summary of Updates

Figures revised; authors added and order of authors updated; Supplemental files updated.

