## Supplementary Figures for "The histone chaperone CAF-1 prevents Rad52-mediated instability of the budding yeast ribosomal DNA during replication-coupled DNA double-strand break repair"

**Figure S1**

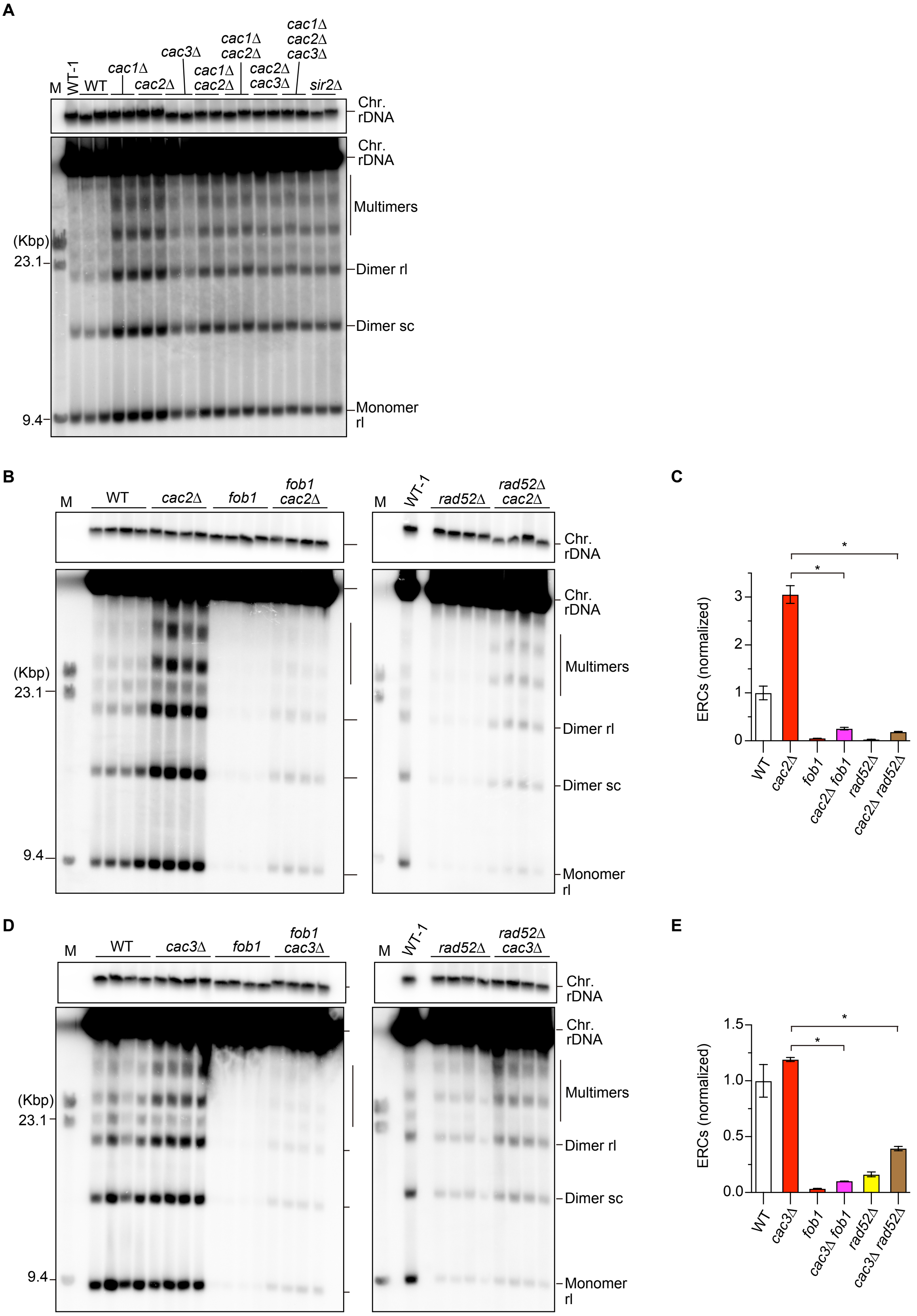


**Supplementary Figure 1. ERC detection in the *caf-1* mutants.**

(A, B, D) ERC detection. DNA was isolated from the two independent clones (A) and four independent clones (B, D) of the indicated strains and separated by agarose gel electrophoresis, followed by Southern blotting with the probe 1, as shown in Fig. 1A. Chromosomal rDNA and different forms of ERCs are indicated. rl and sc indicate relaxed and supercoiled ERCs, respectively. Supercoiled monomers ran off from the gel in the electrophoresis condition under these conditions. The sizes of lambda Hind III DNA markers are indicated. The top panel shows a short exposure of chromosomal rDNA signals. The DNA sample that was indicated as WT-1 was the same DNA sample isolated from the clone that was loaded a separate gel: WT-1 in (A) is the first WT clone in Fig. 1C; WT-1 in (B) is the first WT clone on the left gel; WT-1 in (D) is the first WT clone on the left gel. These DNA samples were used to normalize ERC levels in the mutants that were analyzed on separate gels. (C, E) Quantitation of ERCs. In (C) and (E), ERCs were quantified from (B) and (D), respectively. Multiple comparisons were performed by one-way ANOVA, followed by Tukey's multiple comparisons test; *, statistically significant difference ($p < 0.05$).

**Figure S2**

**A**

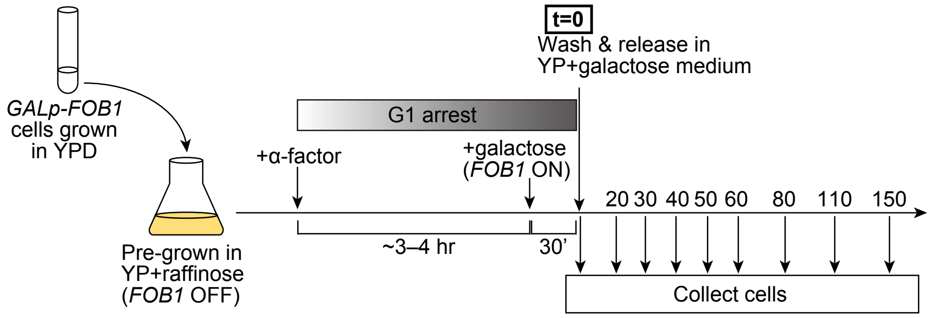


**B**

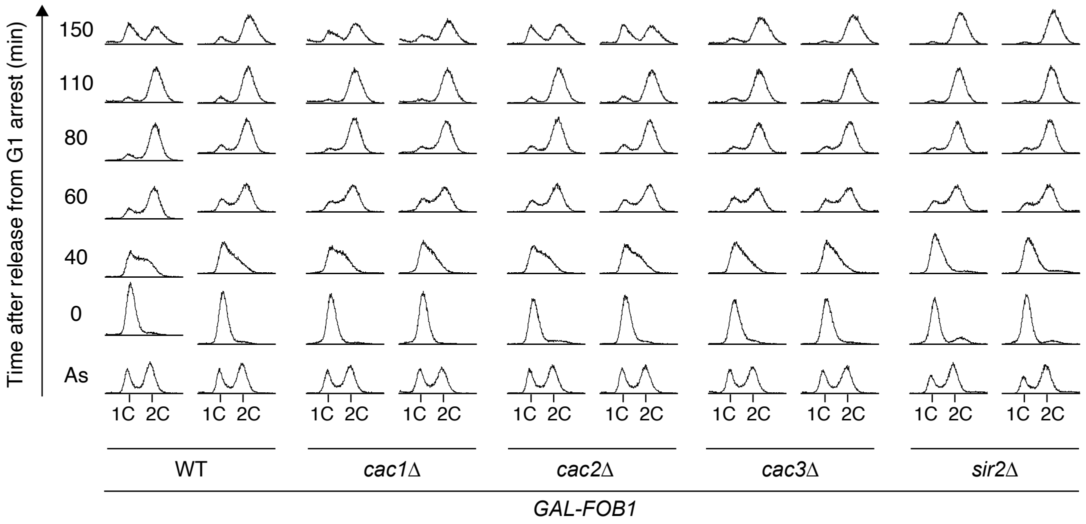


**Supplementary Figure 2. Time course experiments to analyze formation and repair of DSBs.**
(A) Outline of time course experiments.
(B) Cell cycle analysis of two representative time course experiments using GAL-FOB1 WT, *cac1*Δ, *cac2*Δ, *cac3*Δ, and *sir2*Δ strains by flow cytometry.

**Figure S3**

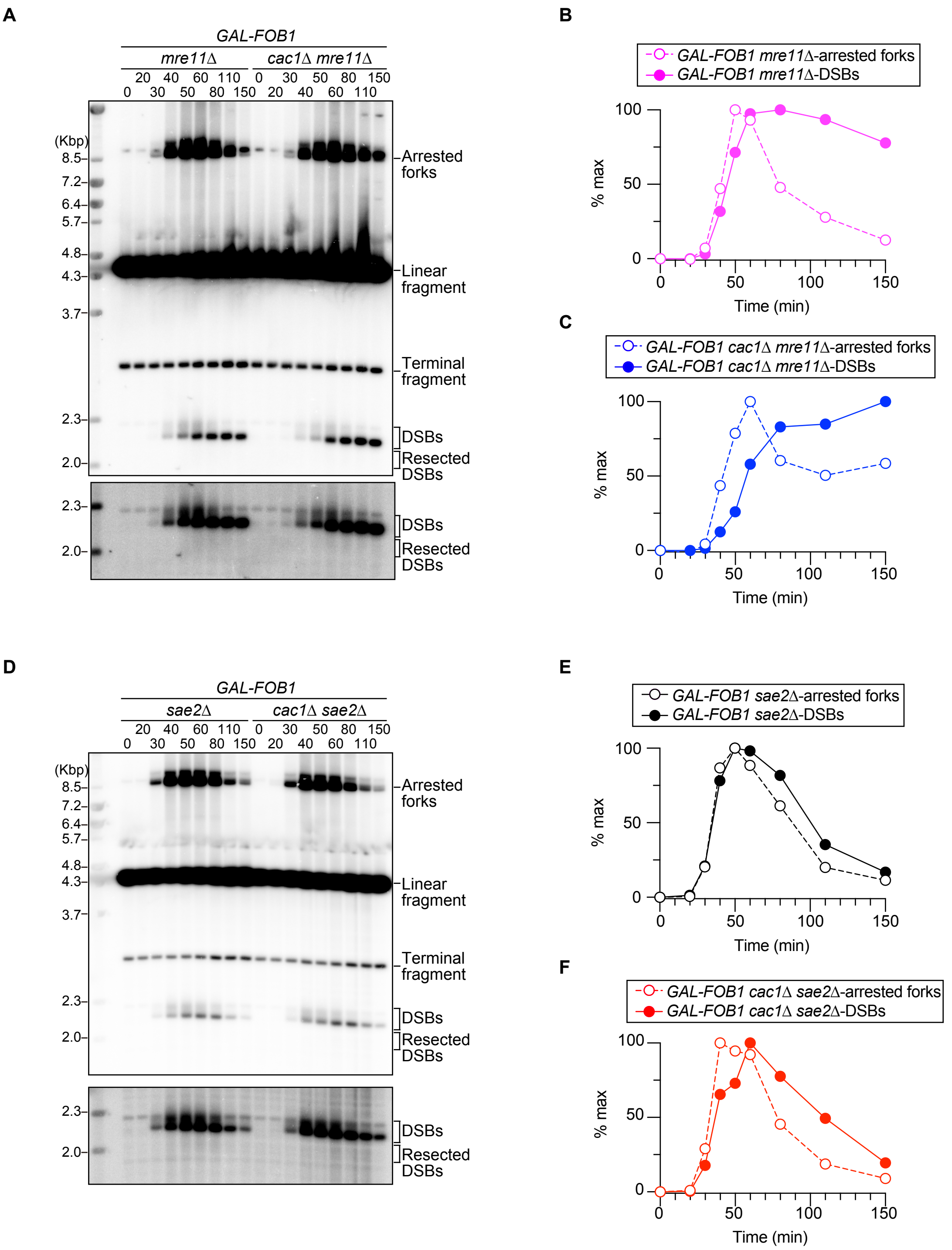


**Supplementary Figure 3. Time course experiments to analyze DSB repair in *cac1*Δ *mre11*Δ and *cac1*Δ *sae2*Δ mutants.** (A, D) DSB assay. Time-course experiments were conducted, as described in Supplementary Figure S2A. Genomic DNA was isolated, digested with Bgl II, separated by size, and analyzed by Southern blotting with probe 4, as shown in Figure 3D. Arrested forks, linear fragments, DSBs and resected DSBs are indicated. Terminal fragments indicate the telomere-proximal rDNA repeat. Long exposure panels around DSBs and resected DSBs are shown below.

(B, C, E, F) Arrested forks and DSBs are expressed as percent of maximum values in one of the two independent time course experiments of the indicated strains.

**Figure S4**

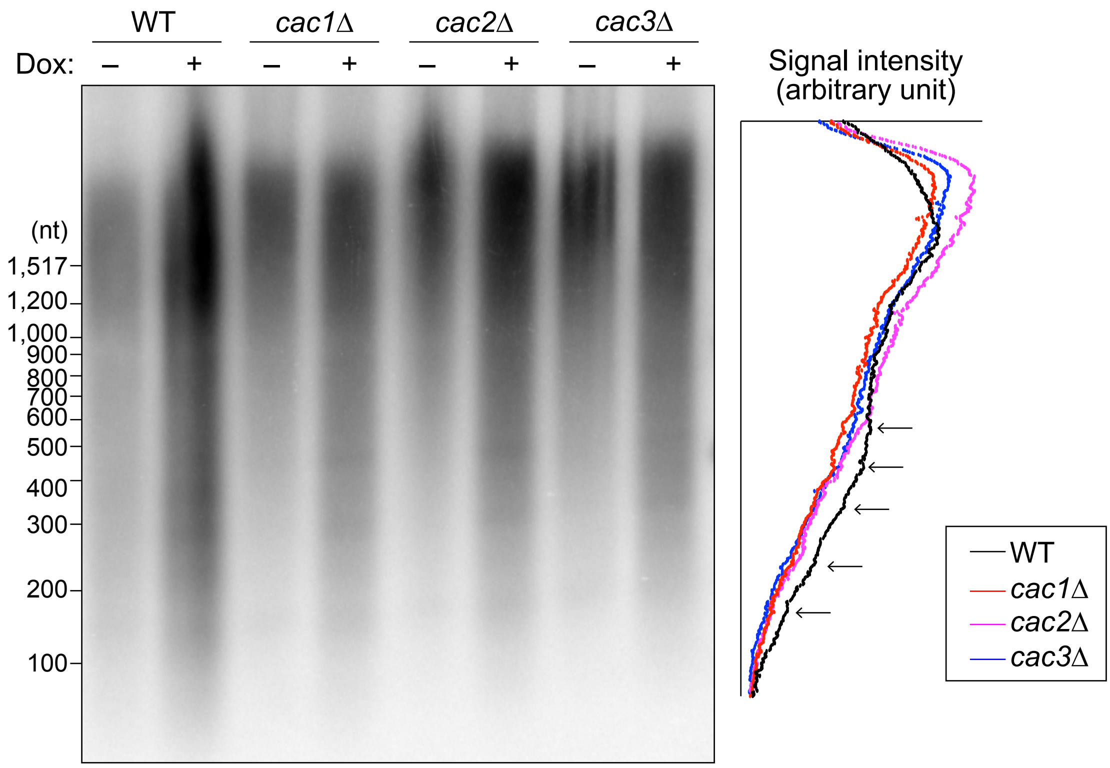


**Supplementary Figure 4. The size of Okazaki fragments in the *caf-1* mutants.**
Okazaki fragments were analyzed in WT, *cac1*Δ, *cac2*Δ, and *cac3*Δ cells carrying a doxycycline-repressible allele of *CDC9* that were treated or not treated with doxycycline (Dox). Genomic DNA was isolated and separated on a denaturing agarose gel, followed by Southern blotting with probe 1, as indicated in Fig. 1A. Lane profiles of the indicated strains, in which *CDC9* was shut off, were included. The positions of the bands of Okazaki fragments are indicated by arrowheads.
