## Supplementary Table for "The histone chaperone CAF-1 prevents Rad52-mediated instability of the budding yeast ribosomal DNA during replication-coupled DNA double-strand break repair"

**Supplementary Table 1. *S. cerevisiae* strains used in this study**

| Name | Genotype |
|---|---|
| HFY3 | *MATa/α, cac1Δ::kanMX/CAC1, fob1::LEU2/FOB1, rad52Δ::hphMX/RAD52* |
| HFY4 | *MATa/α, cac2Δ::kanMX/CAC2, fob1::LEU2/FOB1, rad52Δ::hphMX/RAD52* |
| HFY5 | *MATa/α, cac3Δ::kanMX/CAC3, fob1::LEU2/FOB1, rad52Δ::hphMX/RAD52* |
| HFY18 | *MATa/α, pol30Δ::kanMX/POL30, pBL230-0* |
| HFY19 | *MATa/α, pol30Δ::kanMX/POL30, pBL230-6* |
| HFY20 | *MATa/α, pol30Δ::kanMX/POL30, pBL230-8* |
| HFY21 | *MATa/α, pol30Δ::kanMX/POL30, pBL230-79* |
| HFY32 | *MATa, cac1Δ::hphMX* |
| HFY33 | *MATa, cac2Δ::hphMX* |
| HFY34 | *MATa, cac3Δ::hphMX* |
| HFY70 | *MATa, E-proΔ::GAL1/10-URA3* |
| HFY73 | *MATa, E-proΔ::GAL1/10-URA3, cac1Δ::kanMX* |
| HFY76 | *MATa, E-proΔ::GAL1/10-URA3, sir2Δ::kanMX* |
| HFY106 | *MATa, MCD1-6H10FLAG::kanMX, sir2Δ::URA3* |
| HFY122 | *MATa, MCD1-6H10FLAG::kanMX, cac1Δ::hphMX* |
| MSY45 | *MATa* |
| MSY360 | *MATa, NatNT2-GALL-FOB1, bar1::LEU2* |
| MSY426 | *MATa, MCD1-6H10FLAG::kanMX* |
| MSY613 | *MATa, cdc9::tetO7-CDC9, cmv_LacI-natr, bar1::LEU2* |
| MSY920 | *MATa, sir2Δ::kanMX* |
| MSY937 | *MATa, NatNT2-GALL-FOB1, bar1::LEU2, sir2Δ::hphMX, hmlΔ::kanMX* |
| MSY1231 | *MATa, NatNT2-GALL-FOB1, bar1::LEU2, cac2Δ::kanMX* |
| MSY1261 | *MATa, NatNT2-GALL-FOB1, bar1::LEU2, cac1Δ::kanMX* |
| MSY1262 | *MATa, NatNT2-GALL-FOB1, bar1::LEU2, cac3Δ::kanMX* |
| MSY1376 | *MATa/α, cac1Δ::hphMX/CAC1, YCp22* |
| MSY1377 | *MATa/α, cac1Δ::hphMX/CAC1, YCp22-CAC1* |
| MSY1380 | *MATa/α, cac1Δ::hphMX/CAC1, YCp22-cac1-20* |
| MSY1445 | *MATa/α,, NatNT2-GALL-FOB1/FOB1, bar1::LEU2/BAR1,* |
| MSY1446 | *MATa/α,, NatNT2-GALL-FOB1/FOB1, bar1::LEU2/BAR1,* |
| MSY1450 | *MATa, E-proΔ::GAL1/10-URA3, cac1Δ::kanMX* |
| MSY1453 | *MATa, E-proΔ::GAL1/10-URA3, sir2Δ::kanMX* |
| MSY1550 | *MATa/α, cac1Δ::hphMX/CAC1, cac2Δ::kanMX/CAC2,* |
| MSY1599 | *MATa, cdc9::tetO7-CDC9, cmv_LacI-natr, bar1::LEU2, cac1Δ::hphMX* |
| MSY1604 | *MATa, cdc9::tetO7-CDC9, cmv_LacI-natr, bar1::LEU2, cac2Δ::hphMX* |
| MSY1605 | *MATa, cdc9::tetO7-CDC9, cmv_LacI-natr, bar1::LEU2, cac3Δ::hphMX* |
| MSY1802 | *MATa/α, cac2Δ::kanMX/CAC2, mre11Δ::klTRP/MRE11* |
| MSY1813 | *MATa/α, cac2Δ::kanMX/CAC2, exo1Δ::klTRP/EXO1, sgs1Δ::hphMX/SGS1* |
| MSY1886 | *MATa, cac1Δ::hphMX, YCp22* |
| MSY1890 | *MATa, cac1Δ::hphMX, YCp22-CAC1-9myc* |
| MSY1894 | *MATa, cac1Δ::hphMX, YCp22-cac1-20-9myc* |
| TAK2004a | *MATa, E-proΔ::GAL1/10-URA3* |

All strains are derivatives of W303, which is *ade2-1, ura3-1, his3-11, 15, trp1-1, leu2-3, 112, can1-100*, *rad5*. MSY613 was a *bar1::LEU2* derivative of yIW310, which is a kind gift of Iestyn Whitehouse.
